# Mechanistic basis of teichoic acid transport by a gatekeeper flippase

**DOI:** 10.64898/2026.02.22.707263

**Authors:** Gonzalo Cebrero, Amrutha H. Chidananda, Eric Cester, Julien Dénéréaz, Elif Sena Demir, Alen T. Mathew, D. Ryan Bhowmik, Mario de Capitani, Jean-Louis Reymond, Natarajan Kannan, Fikri Y Avci, Jan-Willem Veening, Ahmad Reza Mehdipour, Camilo Perez

## Abstract

The cell wall is a complex structure that protects bacteria from environmental threats. Phosphocholine-containing teichoic acids are key cell wall biopolymers critical for host colonization, immune evasion, competence, and persistence in *Streptococcus pneumoniae*. The flippase TacF, a member of the multidrug/oligosaccharide-lipid/polysaccharide (MOP) superfamily, monitors the phosphocholine content of teichoic acids during transport, yet the underlying mechanism of this process remains unknown. We present a cryo-EM structure of *S. pneumoniae* TacF in lipid nanodiscs. *In vivo* complementation assays and molecular dynamics simulations reveal key residues involved in teichoic acid recognition and transport, while coevolutionary and conservation analyses delineate common mechanistic elements among MOP flippases, indicating a shared mechanism for polyprenyl-diphosphate-linked oligosaccharide lipid transport. Our findings provide mechanistic insights into an essential flippase involved in *S. pneumoniae* pathogenesis and a potential drug target.

---

Bacteria regulate their cell wall composition through highly coordinated mechanisms crucial for their survival, growth, and adaptability to environmental stresses, such as changes in pH, temperature, or exposure to antibiotics^1–5^. Controlled modulation of the recycling, abundance, and composition of cell wall biopolymers, such as peptidoglycan and teichoic acids, allows pathogens to evade host immune defenses and establish infection^1–4,6^. Dysregulation of these processes can lead to cell lysis or growth arrest, making the different pathways and proteins involved in cell wall content regulation, primary targets of many antibiotics and a key avenue for therapeutic intervention^4,7–10^.

Teichoic acids are essential components of the cell wall of Gram-positive bacteria, constituting about 50% of the cell wall mass^2,11^. In *S. pneumoniae*, teichoic acids play a key role in maintaining cell wall structure, regulating cell division, and contributing to pathogenicity^1,12–14^. They extend from the cell wall surface to facilitate adherence to host cells, allowing bacteria to colonize different types of host tissues^15,16^, and play essential roles in ion homeostasis and antimicrobial resistance^17–19^. *S. pneumoniae* wall teichoic acids (WTA), covalently attached to the peptidoglycan layer, and lipoteichoic acids (LTA), anchored to the cytoplasmic membrane, are composed of repeating units consisting of 2-acetamido-4-amino-2,4,6-trideoxygalactose, glucose (Glc), ribitol phosphate, and two N-acetylgalactosamine (GalNAc) residues modified with phosphocholine groups^20–23^ (**Figure 1A**). Modification of teichoic acids with phosphocholine is critical for *S. pneumoniae* virulence^12,14,24–26^. Disruption of the teichoic acid synthesis pathway^25,27–29^ or exchange of choline for other amino alcohols^25,30^ leads to defective cell division, impaired autolysis, and loss of transformation capacity^27^. The primary importance of phosphocholine lies in its role as an anchoring molecule for a superfamily of cell wall-associated choline-binding proteins (CBPs). These include, among others, LytA^31^, PspA^32,33^, and CbpA^34^, which contribute to evasion of host immune responses^35–37^, attachment to host cells^36,38^, and promote the release of toxins that damage host tissue^31–34,38,39^.

**Figure 1.**
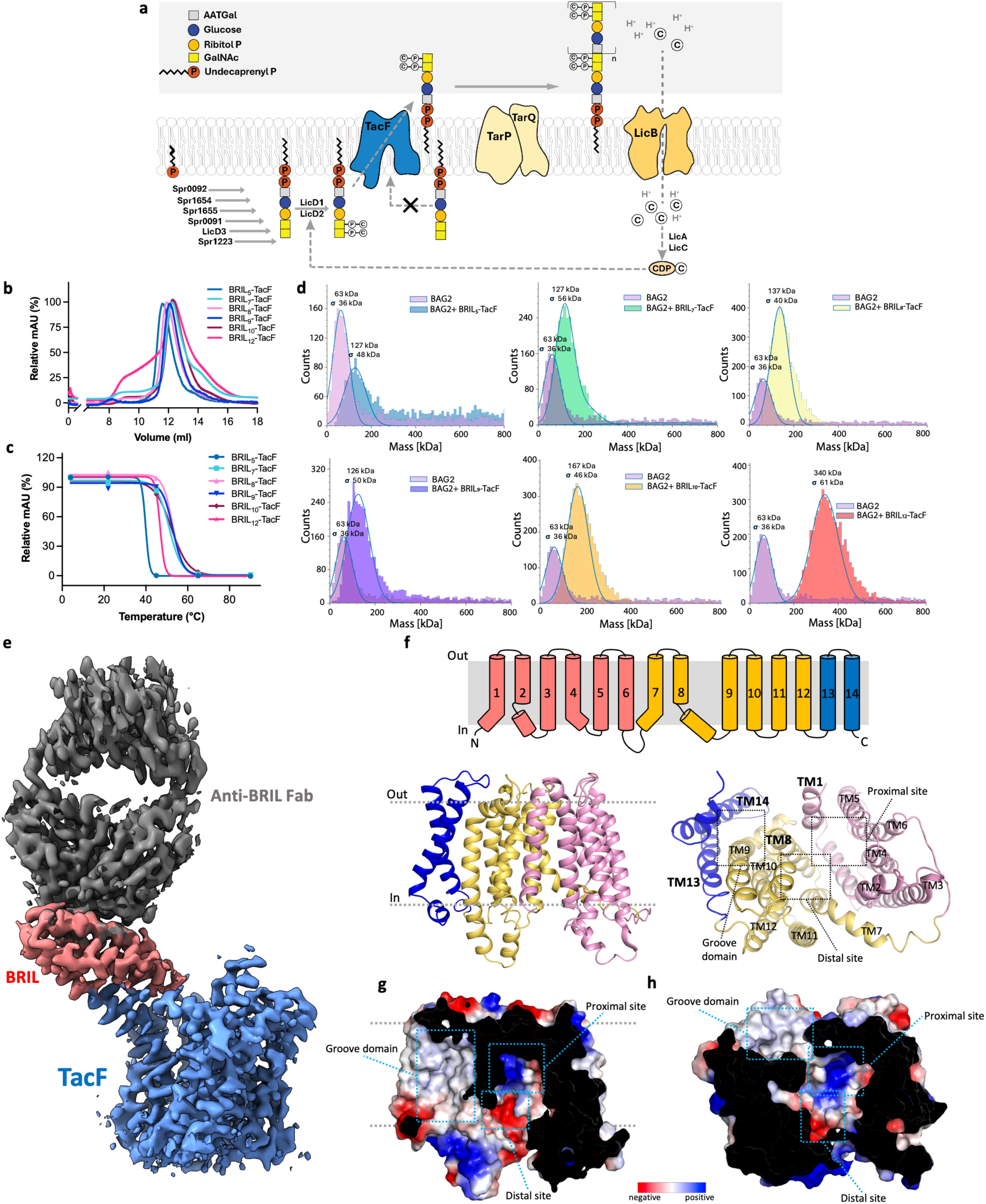
Cryo-EM structure of TacF in lipid nanodiscs. **A**. Teichoic acid synthesis pathway in *S. pneumoniae*. A teichoic acid repeating unit with phosphocholine modifications is assembled on the cytoplasmic side of the membrane and then translocated across it by TacF. The complete teichoic acid is fully polymerized by TarP and TarQ on the extracellular side. **B**. Size exclusion chromatography profiles of BRIL-TacF constructs. **C**. Thermostability assay of purified BRIL-TacF constructs. **D**. Mass photometry analysis of the interaction between BAG2 (Fab) and BRIL-TacF constructs. The species at around 60 kDa corresponds to BAG2 (MW = 52.0 kDa), whereas the species at around 130 kDa corresponds to a 1:1 complex of BRIL-TacF:BAG2 (MW = 134.5 kDa). **E**. Cryo-EM reconstruction map of BRIL_9_-TacF (red and blue, respectively) bound to BAG2 (grey) at 3.63 Å. **F**. Topology of TacF highlighting the N-terminal domain (TM1 to 6, pink), the C-terminal domain (TM7 to 12, yellow), and the external helices TM13 and 14. **G a**nd **H**. Side and cytoplasmic views of a surface electrostatic potential representation of TacF, showing the location of the proximal and distal sites, as well as the groove domain.

*TacF* (*spr1150*) is an essential gene for *S. pneumoniae*, as it monitors the phosphocholine content of teichoic acids by an unknown mechanism, preventing the transport of unmodified assemblies^40–42^ (**Figure 1A**). Following translocation, teichoic acids are polymerized on the outside of the membrane by TarP/TarQ, which are topologically similar to Wzy/Wzz proteins^43^ (**Figure 1A**). The direct contribution of TacF to regulating the cell wall phosphocholine content and the importance of this modification for the localization and function of CBPs make TacF a highly attractive drug target. Deletion of the *tacF* gene is lethal for *S. pneumoniae*^40,42^.

TacF belongs to the multidrug exporter/oligosaccharidyl-lipid/polysaccharide (MOP) superfamily of transporters^23,40,42^, which includes the lipid-II flippase MurJ involved in the biosynthesis of peptidoglycan^44^, the lipid-linked oligosaccharide flippase RFT1 involved in protein N-glycosylation^45^, and the lipid-linked oligosaccharide flippase Wzx involved in O-antigen biosynthesis^46^. However, mechanistic insights into how flippases of this superfamily recognize and translocate their lipid substrates have been limited to MurJ^47–51^. No mechanistic studies have been reported for a teichoic acid flippase within the MOP superfamily, leaving a significant gap in our understanding of their contribution to cell wall assembly in Gram-positive bacteria.

To reveal the mechanism of teichoic acid proofreading and flipping by *S. pneumoniae* TacF, we determined its structure by single-particle cryo-Electron Microscopy (cryo-EM) in lipid nanodiscs. In combination with functional in vivo assays, molecular dynamics (MD) simulations, evolutionary coupling, and conservation analysis, we identified key mechanistic elements involved in the recognition and transport of phosphocholine-modified teichoic acid. The analysis of TacF architecture and other MOP flippases suggests that their mechanism for polyprenyl-diphosphate-linked oligosaccharide recognition and transport might be conserved within this superfamily. This study enhances our understanding of teichoic acid flippases and opens new avenues for targeting *S. pneumoniae* cell wall biosynthesis.

## Results

### Design and analysis of TacF fusion proteins for structure determination

Single-particle Cryo-EM analysis of TacF wild-type (WT) reconstituted into nanodiscs revealed a homogeneous particle distribution and 2D classes; however, it did not produce a high-resolution reconstruction map (**Suppl. Fig. 1**)^52^. We generated a TacF construct fused to a fiducial marker “BRIL”, a four-helix bundle domain from apocytochrome b562a^53^, which is recognized by an affinity-matured semi-synthetic antibody fragment (BAG2) that binds BRIL with high affinity^54^. We designed six BRIL-TacF constructs, where BRIL was attached to the N-terminal domain of TacF. A varying number of TacF N-terminal residues were truncated, allowing us to screen for constructs with diverse conformational flexibility (**Figure 1B-D**, **Suppl. Fig. 2**, **and Suppl. Table 1**). By generating predictive models of these constructs using AlphaFold^55^, we evaluated whether: (i) the segment between BRIL and TacF showed a helical secondary structure, decreasing the likelihood of a flexible BRIL construct; (ii) the segment connecting BRIL and TacF displayed an AlphaFold pLDDT confidence score higher than 70; and (iii) the BRIL segment introduced potential folding artifacts. The constructs BRIL_5_-TacF and BRIL_9_-TacF, which lack five and nine residues from the N-terminal domain, respectively, exhibited the most favorable characteristics (**Suppl. Fig. 2A**). We expressed and purified the six BRIL-TacF constructs, and assessed their monodispersity and thermostability (**Figure 1B,C and Suppl. Fig. 2B**). Except for BRIL_12_– TacF, all the other constructs displayed a single monodisperse peak in size-exclusion chromatography (**Figure 1B**). BRIL_5_-TacF and BRIL_12_-TacF exhibited the lowest thermostability (**Figure 1C**), whereas BRIL_7_-, BRIL_8_-, BRIL_9_-, and BRIL_10_-TacF exhibited the highest (**Figure 1C**). To evaluate whether the BRIL-TacF constructs retained the ability to form complexes with the BAG2 anti-BRIL Fab, we performed mass photometry analysis with the purified BRIL-TacF constructs and BAG2, where we expected to see a population around 110-130 kDa corresponding to a 1:1 complex of BRIL-TacF:BAG2 (BRIL-TacF MW = 82.5 kDa, BAG2 MW = 52.0 kDa) (**Figure 1D**). Our results show that incubation of BRIL_5_-, BRIL_7_-, BRIL_8_-, and BRIL_9_-TacF with BAG2 allows the formation of a 1:1 BRIL_i_-TacF:BAG2 complex, whereas incubation of BRIL_10_– and BRIL_12_-TacF with BAG2 yields species of higher mass, likely corresponding to complexes containing multiple BAG2 Fab or to aggregates. Hence, based on the monodispersity of the purified proteins, their thermostability, the capacity to form 1:1 complexes with BAG2, and the analysis of AlphaFold models, we selected BRIL_9_-TacF to pursue single particle cryo-EM analysis.

### Cryo-EM structure of BRIL_9_-TacF in nanodiscs

Purified BRIL_9_-TacF was incorporated in nanodiscs containing a mixture of 1-palmitoyl-2-oleoyl-sn-glycero-3-phospho-(1’-rac-glycerol) (POPG) and diacylglycerol (DAG) (3:1, w/w). POPG and DAG-based glycolipids are major components of the *S. pneumoniae* membrane^56,57^. The nanodiscs sample was incubated with BAG2 antibody and further purified using size-exclusion chromatography (**Suppl. Fig. 3A**)^52^. Iterative 2D classification resulted in clear features characteristic of the BAG2 Fab bound to BRIL and secondary structural features of TacF embedded in nanodiscs (**Suppl. Fig. 3B**). Further particle sorting, masking, and 3D refinement leading to an BRIL_9_-TacF:BAG2 EM map reconstructed to an overall 3.63 Å resolution, with the TacF part displaying a resolution ranging from 2.5 Å to 3.5 Å (**Figure 1E, Suppl. Fig. 3C-E, 4, and Table 1**). The resulting map showed clear density for the whole TacF protein except for nine residues from the first cytoplasmic loop (residues 68-76). The interaction between the BRIL segment and the BAG2 Fab was mediated by both the light and heavy chains of BAG2, involving extensive Van der Waals, polar, and cation-π interactions of its complementarity-determining regions (CDRs) and helices 2 and 3 of BRIL (**Suppl. Fig. 5A**).

**Table 1.** Cryo-EM data collection, refinement and validation statistics.

|  | **BRIL<sub>9</sub>-TacF:BAG2 in nanodiscs**<br>**(PDB ID 9XZN, EMD-72367)** |
| --- | --- |
| **Data collection and processing** |  |
| Microscope | Glacios TEM |
| Camera | Gatan K3 GIF |
| Magnification | 46,000 |
| Voltage (kV) | 200 |
| Electron exposure (e<sup>-</sup>/Å<sup>2</sup>) | 55 |
| Defocus range (μm) | -0.5 to -3 |
| Pixel size (Å) | 0.878 |
| Symmetry imposed | C1 |
| Initial particle images (no.) | 6,591 |
| Final particle images (no.) | 53,717 |
| Map resolution (Å) | 3.63 |
| FSC threshold | 0.143 |
| Map resolution range (Å) | 2.5-4.0 |
| **Refinement** |  |
| Initial model used | AlphaFold |
| Model resolution (Å) | 3.8 |
| FSC threshold | 0.5 |
| Map sharpening b-factor (Å<sup>2</sup>) | -126 |
| EMRinger score (BRIL<sub>9</sub>-TacF:BAG2) | 1.22 |
| **Model composition** |  |
| Non-hydrogen atoms | 4668 |
| Protein residues | 580 |
| **B-factors (Å<sup>2</sup>)** |  |
| Protein | 53.30 |
| **R.m.s. deviations** |  |
| Bond lengths (Å) | 0.002 |
| Bond angles (°) | 0.503 |
| **Validation** |  |
| MolProbity score | 1.71 |
| Clashscore | 6.53 |
| Poor rotamers (%) | 1.17 |
| **Ramachandran plot** |  |
| Favored (%) | 95.67 |
| Allowed (%) | 4.15 |
| Disallowed (%) | 0.17 |

FSC, Fourier Shell Correlation

TacF displays a fold comprised of 14 transmembrane (TM) helices organized into three distinct domains, an N-terminal domain (TM1-6), a C-terminal domain (TM7-12), and a two-TM domain (TM13-14) that contributes to the formation of a hydrophobic groove external to the transporter core, ‘groove domain’ (**Figure 1F-H**)^52^. The N– and C-terminal domains exhibit a two-fold pseudo-rotational symmetry with only few significant differences on their conformation, leading to an r.m.s.d. of 2.0 Å upon C-alpha superposition (**Suppl. Fig. 5B**). The second helix of each of the N-and C-terminal domains contains an unwound segment on the cytoplasmic side, featuring two motifs, xxG(I/V)xxYG in TM2 and xxVxxPRx in TM8, which are highly conserved among Gram-positive bacterial homologues with over 40% sequence identity (**Suppl. Fig. 5C**). The presence of glycine and proline residues in these motifs contributes to the disruption of the α-helical structure of these TM segments.

The TacF structure exhibits an inward-facing conformation with an entrance flanked by one TM helix from the N-terminal domain and one from the C-terminal domain, TM1 and TM8, positioned adjacent to the groove domain (**Figure 1F**)^52^. The central cavity, surrounded by TM1-2 and TM4-6 from the N-terminal domain, and TM7-8 and TM10-12 from the C-terminal domain, features a large vestibule with a pronounced positive electrostatic surface potential close to the groove domain (proximal site), and a negatively charged region farther from it (distal site) (**Figure 1G-H**). The main residues contributing to the positive surface of the proximal site are R152, R230, R250, and R333, whereas D340 and D394 are the main contributors to the negative charge of the distal site (**Suppl. Fig. 5D**).

### Conserved features of the central cavity and groove domain in Gram-positive bacteria

We investigated the conservation of residues contributing to the formation of the positively charged proximal site, the negatively charged distal site, and the groove domain. A sequence similarity network (SSN) was generated to visualize the relationships among proteins with high sequence identity to TacF across multiple Gram-positive bacteria. The SSN analysis comprises 810 sequences grouped into five distinct clusters, with each node representing either a single protein or a group of sequences sharing more than 95% identity (**Figure 2**). This analysis yields an arrangement of sequences distributed into five distinct clusters, with each cluster comprising sequences that share a sequence similarity of at least 40% (**Figure 2**). Cluster I comprises a large number of sequences from diverse *Streptococcus* species and includes the sequence of *S. pneumoniae* TacF (black dot), along with closely related homologues from *S. infantis*, *S. mitis*, *S. oralis*, *S. peroris*, *S. australis*, *S. gwangjuense*, *S. pseudoneumoniae*, and *S. symci* (**Figure 2**). In contrast, TacF homologues from other *Streptococcus* species, including *S. parasanguinis*, *S. rubneri*, *S. salivarus*, *S. thermophilus*, and *S. vestibularis*, form a separate group (cluster-II), indicating substantial divergence between the TacF proteins of these two clusters. A detailed comparison of the three key functional domains, the groove domain, proximal site, and distal site, revealed that although the electrostatic surface potentials are broadly similar across clusters, the sequence similarity of residues within these domains is significantly higher in cluster II (**Figure 2**). Despite shared surface charge features, the higher sequence conservation in cluster II may reflect functional specialization or evolutionary constraints not present in the more heterogeneous cluster I. When comparing their teichoic acid composition, several *Streptococcus* species in cluster I have been reported to have phosphocholine-decorated teichoic acids^23,58,59^. In contrast, there are no reports of this modification for species in cluster II^60–62^.

**Figure 2.**
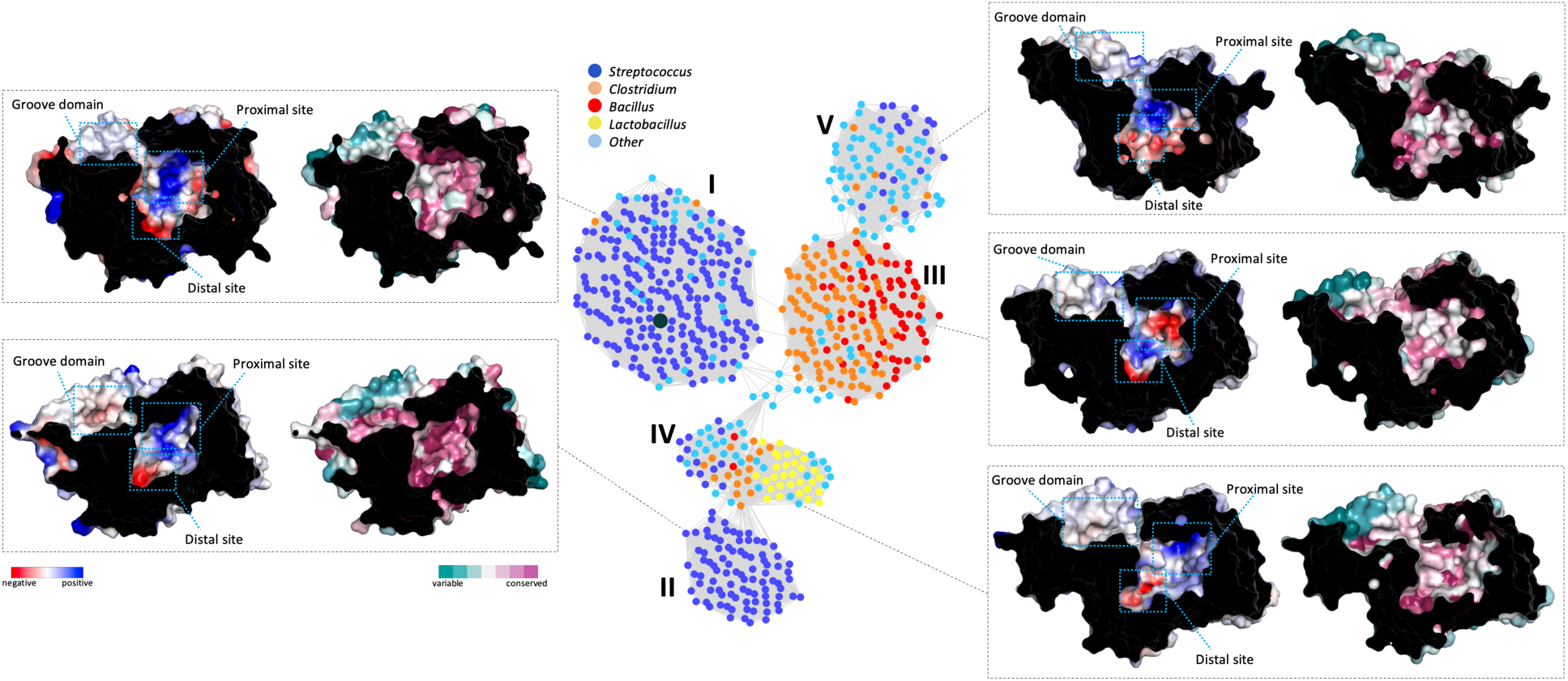
Conservation of TacF and its architecture across Gram-positive bacteria. SSN analysis of TacF homologues from Gram-positive bacteria reveals five distinct clusters. Each node represents a group of sequences sharing >95% identity. Edges between nodes indicate an identity of at least 40% among them. For each cluster, a surface electrostatic potential representation of a representative TacF homologue is shown, along with a sequence conservation analysis for the entire cluster. *S. pneumoniae* TacF, black dot in Cluster-I.

Additionally, we found that sequences from various *Bacillus* and *Clostridium* species form a distinct group (cluster-III), which differs from the *S. pneumoniae* TacF cluster in terms of the electrostatic surface potential shown at the proximal and distal sites (**Figure 2**), although the sequence conservation of residues in the proximal site is low. These contrasting properties of the central cavity may reflect adaptations to the distinct teichoic acid compositions in *Bacillus*, *Clostridium*, and *Streptococcus* species^23,63–65^. On the other hand, cluster-IV and cluster-V, which include a greater diversity of species, display electrostatic surface properties at the proximal and distal sites similar to those of *S. pneumoniae* TacF (**Figure 2**).

Taken together, these results indicate that the hydrophobic properties of the groove domain are broadly conserved among TacF homologues. However, the cluster-specific variations in sequence conservation and the charges of the proximal and distal sites likely reflect specific adaptations to different teichoic acid substrates among Gram-positive bacteria.

### Model of teichoic acid recognition by TacF

We performed Molecular dynamics (MD) simulations to better understand and reveal the interactions of TacF with the teichoic acid molecule (**Figure 3A and Suppl. Fig. 6A,B**). We conducted four independent simulation runs, each lasting at least 500 ns. We initially positioned a teichoic acid molecule near the lateral opening between TM1 and TM8, which moved closer to the entrance during the equilibration phase. In the simulations, the headgroup of the teichoic acid molecule entered the central cavity, with the diphosphate moiety primarily interacting with residues R15 and R269, located in TM1 and TM8, respectively (**Figure 3B,C**). These two helices form the lateral entrance that connects the hydrophobic groove to the central cavity (**Figure 1F**). A similar set of positively charged residues, positioned in TM1 and TM8, was reported to coordinate the diphosphate group of the head group of lipid-II in the flippase MurJ^48,49,66^. In the simulations where the teichoic acid headgroup enters the central cavity, this is mainly stabilized by interactions between the phosphocholine phosphate groups and the two GlcNAc units of the repeating unit with nearby residues at the proximal site (**Figure 3B,D**). Specifically, residues R230 and R333, which are the primary contributors to the positive charge of the proximal site, along with Q249, R250, and T253, contribute to stabilize the teichoic acid headgroup in the central cavity (**Figure 3D**). In contrast, residues from the distal site do not interact with the teichoic acid headgroup.

**Figure 3.**
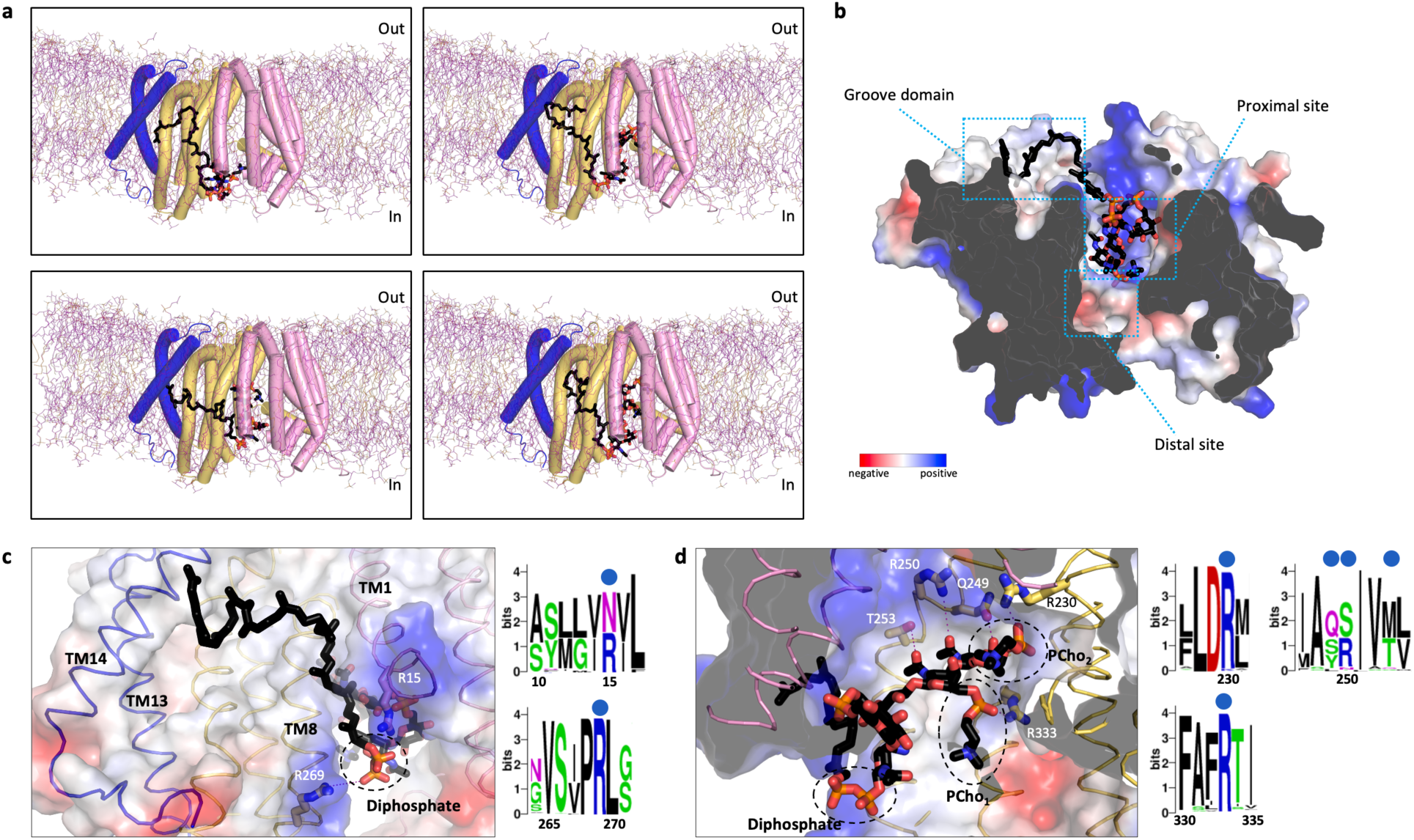
Molecular dynamics simulations of TacF and teichoic acid interaction. **A**. Snapshots from four independent MD simulations of TacF embedded in a heterogeneous bilayer, with a teichoic acid molecule initially positioned near the lateral entrance between TM1 and TM8. **B**. Surface electrostatic potential representation of TacF viewed from the cytoplasmic side of the membrane. The teichoic acid molecule binding is stabilized by interactions of the undecaprenyl tail, diphosphate group, and repeating unit with the groove domain, positively charged residues at the entrance between TM1 and TM8, and the proximal site, respectively. **C**. Interactions of the undecaprenyl tail and diphosphate group. *Right*, sequence logos analysis of the regions containing residues that coordinate the diphosphate group (blue dots). **D**. Interactions of the teichoic acid repeating unit. *Right*, sequence logos analysis of the regions containing residues that coordinate the GlcNAc and phosphocholine units (blue dots). PCho indicates phosphocholine.

The positively charged residues that stabilize the teichoic acid headgroup are highly conserved among TacF homologues found in Gram-positive bacteria, while Q249 and T253 are less conserved (**Figure 3C,D**). We also observed that the undecaprenyl aliphatic moiety of the teichoic acid remained positioned in the groove domain throughout the simulation (**Figure 3A-C and Suppl. Fig. 6B**). Notably, even after extending the simulations by an additional 400 ns, the undecaprenyl chain remained in this location. These results align with previous observations regarding the recognition of the undecaprenyl moiety of lipid-II by the flippase MurJ^48,49,66^, where this moiety was observed interacting with the groove domain.

Together, these results support a model in which TacF recognizes the teichoic acid molecule through four key interactions namely: (i) the undecaprenyl tail is stabilized at the groove domain; (ii) the diphosphate linker is coordinated by residues R15 and R269 at the entrance to the central cavity formed by TM1 and TM8; (iii) the two phosphocholine moieties are stabilized by positively charged residues R230 and R333 at the proximal site through interactions with their phosphate groups; and (iv) the GlcNAc units engage in interactions with Q249, R250 and T253.

### Functional characterization of residues involved in teichoic acid recognition

To assess the significance of residues involved in the teichoic acid headgroup recognition, we performed an in vivo functional complementation assay in the *S. pneumoniae* D39V strain^67^. A *tacF* knock-out strain was generated in a strain harboring two different constructs, each using distinct inducible promoters. The first one, induced by Isopropyl ß-D-1-thiogalactopyranoside (IPTG) via the P*lac* promoter, contained a copy of the wild-type *tacF* gene, while the other, induced by anhydrotetracycline (aTc) via the P*tet* promoter, carried a specific TacF mutant to be tested in the assay. We first complemented the wild type allele of *tacF* by inserting an ectopic copy under an IPTG inducible promoter (Plac) at the ZIP locus^67^. We then replaced the original *tacF* loci with an antibiotic resistance marker to be able to completely titrate the expression of the wild-type allele by IPTG. Each of the identified *tacF* mutants was inserted under an aTc inducible promoter (Ptet) at the *bgaA* locus (**Figure 4A**). This system allowed for a titratable expression of either the wild-type *tacF* allele or a mutant *tacF* allele. To test and verify the double expression system, we inserted a wild-type *tacF* allele under both promoters. Transformed cells were used to inoculate three separate cultures: one with IPTG, another with aTc, and a third without IPTG or aTc. The growth curve comparison suggested that *tacF* deletion is lethal for pneumococcus, and could be complemented by either IPTG or aTc induction (**Figure 4A**).

**Figure 4.**
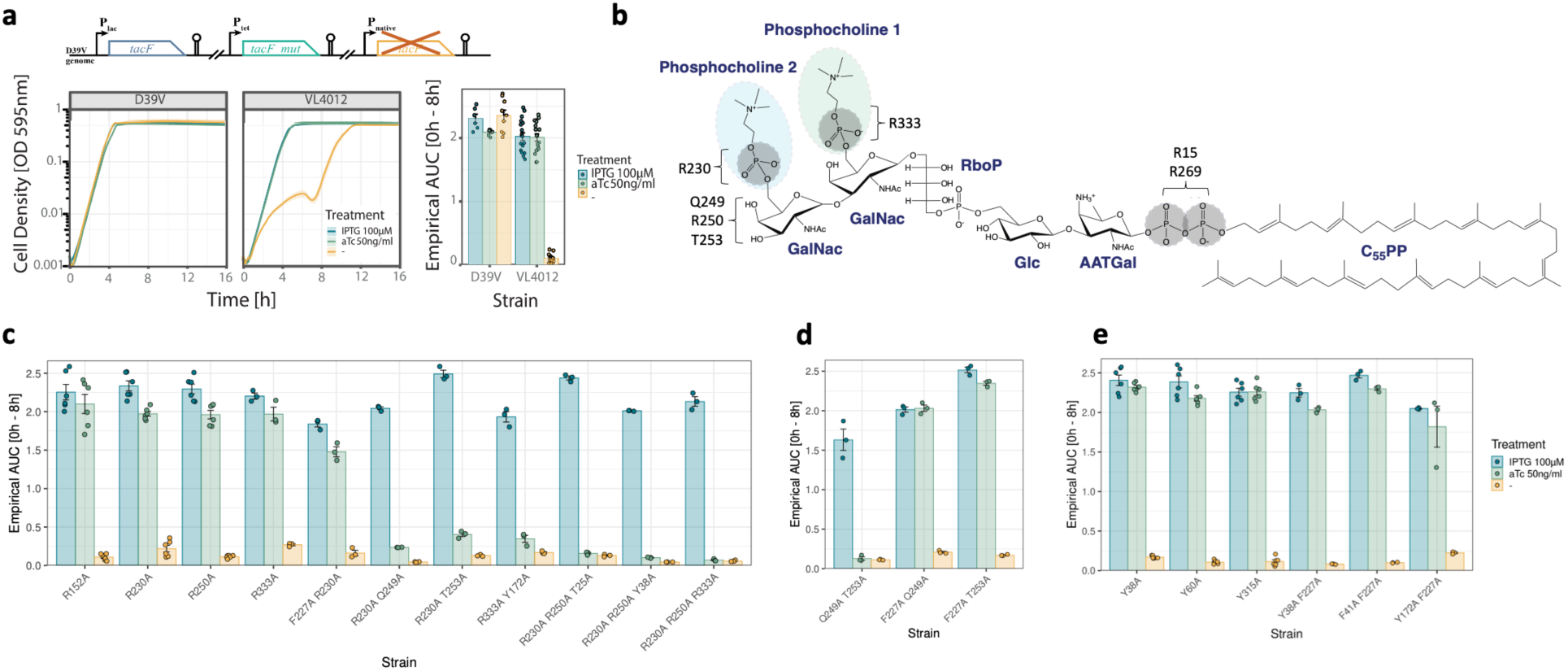
Characterization of the binding partners of the teichoic acid repeating unit. **A**. Double expression system of TacF. A wild type *tacF* allele was inserted under Plac promoter in the ZIP locus. A second *tacF* allele was inserted under Ptet promoter, at the bgaA locus. The native *tacF* allele was replaced with a selection marker. Growth curves of *S. pneumoniae* D39V, and VL4012 carrying the double expression system for the *tacF* wild type allele under both Plac and Ptet promoters. Cultures were grown in media supplemented with IPTG (blue), aTc (green), or no inducer (yellow). Bar plots show the empirical area under the growth curves (AUC) from 0 to 8 hours. Colors represent the same induction conditions as in the growth curves. **B**. Chemical structure of a teichoic acid molecule annotated with key TacF-interacting residues identified from MD simulations. **C-E**. Bar plots showing the growth phenotype of *S. pneumoniae* strains carrying the double expression system for *tacF* variants (see growth curves in **Suppl. Fig. 7**). n = 3 biological replicates; n = 3 technical replicates, for all experiments.

We generated strains carrying alanine substitutions designed to disrupt TacF recognition of phosphocholine moieties and GlcNAc units in the teichoic acid repeating unit (**Figure 4B-D and Suppl. Fig. 7**). Our findings indicate that double and triple mutants of these residues led to substantial growth defects in *S. pneumoniae*, while single mutations had no significant impact on growth. This reflects that recognition of large molecules like teichoic acid depends on cooperation among several binding partners. Taken together, these results suggest that destabilizing the binding of the phosphate groups of the phosphocholine moieties and the GlcNAc units in the teichoic acid headgroup is sufficient to compromise the fitness of *S. pneumoniae*.

Proteins that bind choline frequently coordinate this molecule via cation-π interactions established between aromatic side chains and the positive charge of choline^68–71^. TacF features several aromatic residues in its central cavity (**Suppl. Fig. 8A)**. To study the contribution of these residues to recognizing the teichoic acid repeating unit, we introduced alanine substitutions alone or combined with substitutions of residues involved in coordinating phosphocholine and GlcNAc units, as described above (**Figure 4C-E and Suppl. Fig. 7)**. Our findings indicate that replacing the aromatic residues does not affect *S. pneumoniae* growth, unless they are combined with variants of the positively charged residues that coordinate the phosphate group in the phosphocholine moieties. Further demonstrating the significance of the phosphocholine modification in teichoic acid transport.

### Evolutionary coupling analysis of TacF

We performed evolutionary coupling analysis to identify coevolving residues, thereby highlighting those of potential functional importance independent of the protein structural conformation^72–74^. We analyzed an alignment of 138,448 sequences and identified 650 long-range evolutionary couplings within the 99th percentile. Our analysis reveals that residues R15 and R269, which coordinate the diphosphate group, as shown by MD simulations, are highly evolutionarily conserved (**Figure 5A**). Similarly, residues R230 and R333, which stabilize the phosphate group of the phosphocholine moieties, as well as Q249, R250, and T253, involved in the coordination of GlcNAc, also show strong evolutionary conservation (**Figure 5A**). Multiple residues located at the groove domain are also highly conserved, emphasizing the functional significance of this domain across homologues (**Figure 5A**).

**Figure 5.**
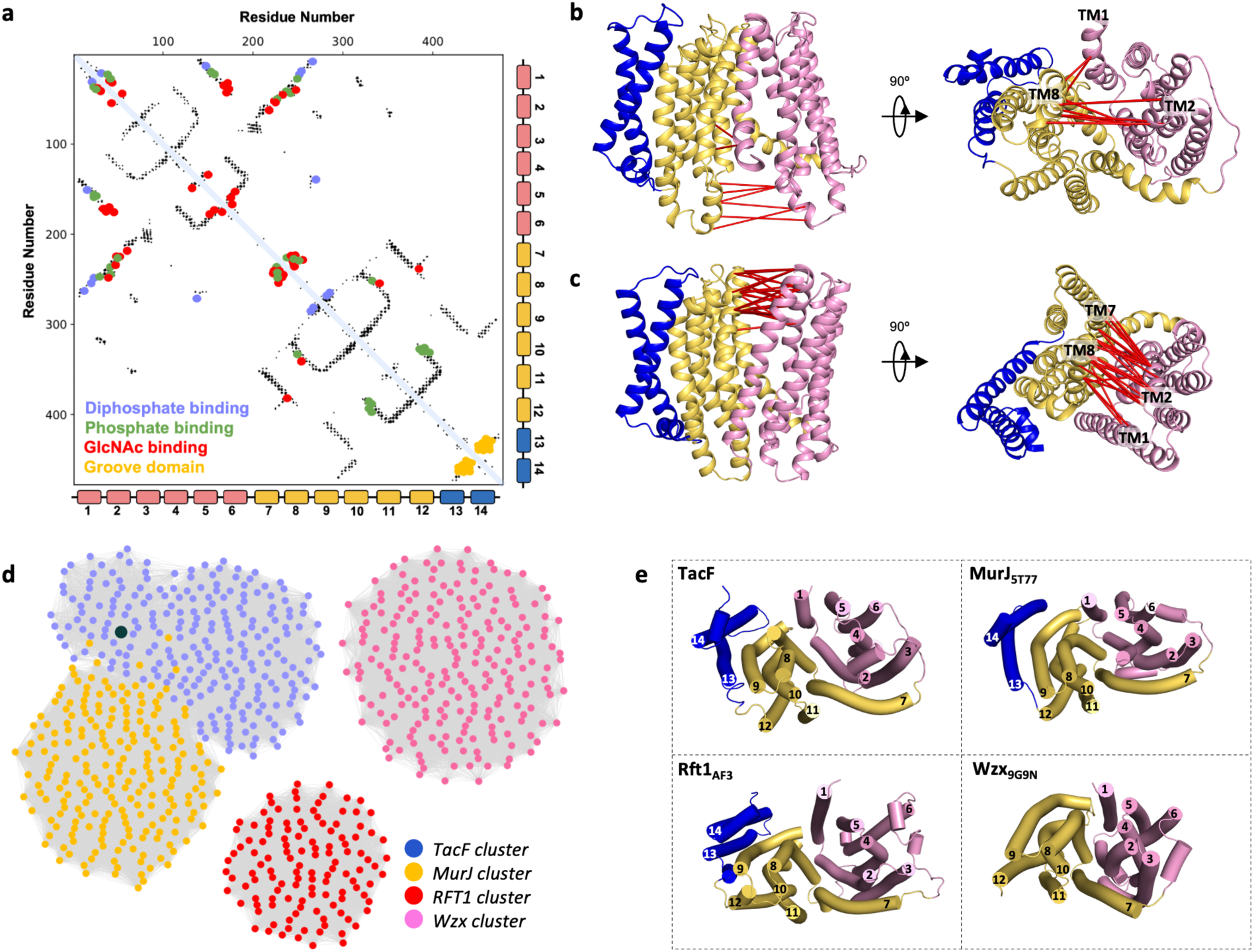
Evolutionary coupling analysis of TacF and SSN analysis of MOP superfamily flippases. **A**. Evolutionary coupling analysis of TacF. Coevolved pairs are shown as black dots. Functionally relevant residues for teichoic acid recognition are shown as overlaid blue dots (diphosphate binding), green dots (phosphate binding), red dots (GlcNAc binding), and yellow dots (groove domain). **B** and **C**. Coevolving pairs (red lines) with long inter-residue distances in the cryo-EM TacF structure (**B**) or in an Alphafold model of TacF in outward conformation (**C**). **D**. SSN analysis of different MOP superfamily flippases. Each node represents a group of sequences sharing >95% identity. Edges between nodes indicate an identity of at least 40% among them. **E**. Comparison of domain distribution and topology of four MOP superfamily flippases viewed from the cytoplasmic side of the membrane. TacF, MurJ (PDB ID: 5T77), Rft1 (AlphaFold model), and Wzx (PDB ID: 9G9N). The N-terminal and C-terminal domains are colored pink and yellow, respectively. External helices, TM13 and 14, are colored blue.

In addition, we identified a subset of strongly coevolving residue pairs that are separated by larger distances, ranging from 15.6 to 25.3 Å (**Figure 5B**). These residues, clustered on the cytoplasmic side of TM helices 1, 2 and 8, are predicted to come into close proximity during the transition to the outward-facing state, as suggested by an AlphaFold model of an outward-open state of TacF (**Figure 5C**). Our analysis also shows coevolving residue pairs within <7.91 Å of each other on the extracellular side of TM1, 2, 7, and 8 in the cryo-EM inward-facing TacF structure, involving residues from both the N– and C-terminal domains (**Suppl. Fig. 9**). These pairs are 17.0 to 20.1 Å apart in the outward-open AlphaFold model of TacF (**Figure 5C**). The extracellular and cytoplasmic pairs of residues form a network of Van der Waals and polar interactions that help stabilize the inward– and outward-facing conformations of TacF.

In summary, our analysis indicates that the residues mediating the teichoic acid molecule binding, as well as those that stabilize the inward– and outward-facing states, are under evolutionary selection pressure in bacterial species carrying the *tacF* gene.

## Discussion

The MOP superfamily consists of four subfamilies: (i) the ubiquitous multi-drug and toxin extrusion (MATE) family; (ii) the prokaryotic polysaccharide transporter (PST) family; (iii) the eukaryotic oligosaccharidyl-lipid flippase (OLF) family, and (iv) the bacterial mouse virulence factor family (MVF)^75^. Flippases in this superfamily include TacF and Wzx, both part of the PST family, with Wzx involved in transporting various undecaprenyl diphosphate-linked oligosaccharides that are important for O-antigen synthesis^76,77^; RFT1, a eukaryotic protein belonging to the OLF family, presumably involved in the transport of a dolichol diphosphate-linked oligosaccharide fundamental for N-linked glycosylation^45,78^; and MurJ, a bacterial MVF flippase that catalyzes the flipping of lipid-II, an undecaprenyl pyrophosphate-MurNAc-pentapeptide-GlcNAc precursor essential for peptidoglycan synthesis^48,78^. A sequence similarity network (SSN) analysis of TacF, Wzx, MurJ, and RFT1 homologues reveals that *S. pneumoniae* TacF clusters with multiple uncharacterized proteins that contain a 14-TM helix topology. Within this cluster, several sequences exhibit high similarity to MurJ proteins from diverse bacterial species, which themselves form a distinct yet closely related cluster (**Figure 5D**). This highlights the architectural similarity between the two flippases, despite their classification into different families within the MOP superfamily (**Figure 5E**). In contrast, eukaryotic RFT1 flippases, which also share a similar 14-TM helix architecture (**Figure 5E**), form a distinct and separate cluster, displaying low sequence conservation with TacF and MurJ proteins (**Figure 5D**). Similarly, Wzx flippases form a distinct cluster, consistent with their different 12-TM helix architecture^79^ (**Figure 5D,E**). Unlike TacF, MurJ, and RFT1, Wzx proteins lack the two additional helices (TM13-14) that contribute to the formation of the groove domain (**Figure 5E**).

Multiple crystal structures of MurJ, captured in distinct inward-facing conformations and an outward-open conformation, have been reported previously^48,49,80,81^. Like TacF, MurJ exhibits a 14-TM helix topology, organized into an N-terminal domain (TM1-6), a C-terminal domain (TM7-12), and two additional helices, TM13 and TM14 (**Figure 5E**). Its central cavity features a strongly cationic proximal site and a polar charged distal site^51^ (**Suppl. Fig. 5E**). Similarly, an AlphaFold model of RFT1 reveals a comparable three-domain distribution, with a defined proximal site, a distal site, and a groove domain (**Suppl. Fig. 5F**). The shared structural architecture of TacF, MurJ, and RFT1, along with the conserved features of the groove domain and the distinct proximal and distal sites, suggests a common evolutionary origin for the recognition and translocation of polyprenyl-diphosphate-linked oligosaccharide lipids by MOP superfamily flippases.

Our results support a model in which TacF recognizes the teichoic acid molecule through specific interactions of its different chemical groups. In this model, the hydrophobic groove domain contributes to the recognition of the undecaprenyl aliphatic chain, thereby increasing the likelihood of interaction between the rest of the molecule and key residues at the lateral entrance and within the central cavity (**Figure 6**). Once a teichoic acid molecule is found in the proximity of the lateral entrance formed by TM1 and TM8, the diphosphate linker establishes electrostatic interactions with residues R15 and R269 (**Figure 6**). Our data indicate that once the complete repeating unit of the teichoic acid enters the central cavity, crucial interactions form with the two phosphocholine moieties via their phosphate groups, and with the GlcNAc units of the repeating unit. Recognition of the phosphocholine moieties through their phosphate groups is a key element in the molecular mechanism by which TacF monitors the phosphocholine content of teichoic acids, preventing the transport of unmodified teichoic acids^40–42^. Our findings on the importance of recognizing phosphocholine moieties through their phosphate groups align with earlier studies on *S. pneumoniae* growth in media where choline is replaced with ethanolamine,^27,82^ since incorporation of phosphoethanolamine into teichoic acids preserves the key elements recognized by TacF in the repeating unit, specifically the phosphate groups and GlcNAc. These findings suggest that the trimethylammonium group of the phosphocholine moieties plays a key role in *S. pneumoniae*, mainly because it anchors CBPs to the cell wall, rather than facilitating recognition by TacF.

**Figure 6.**
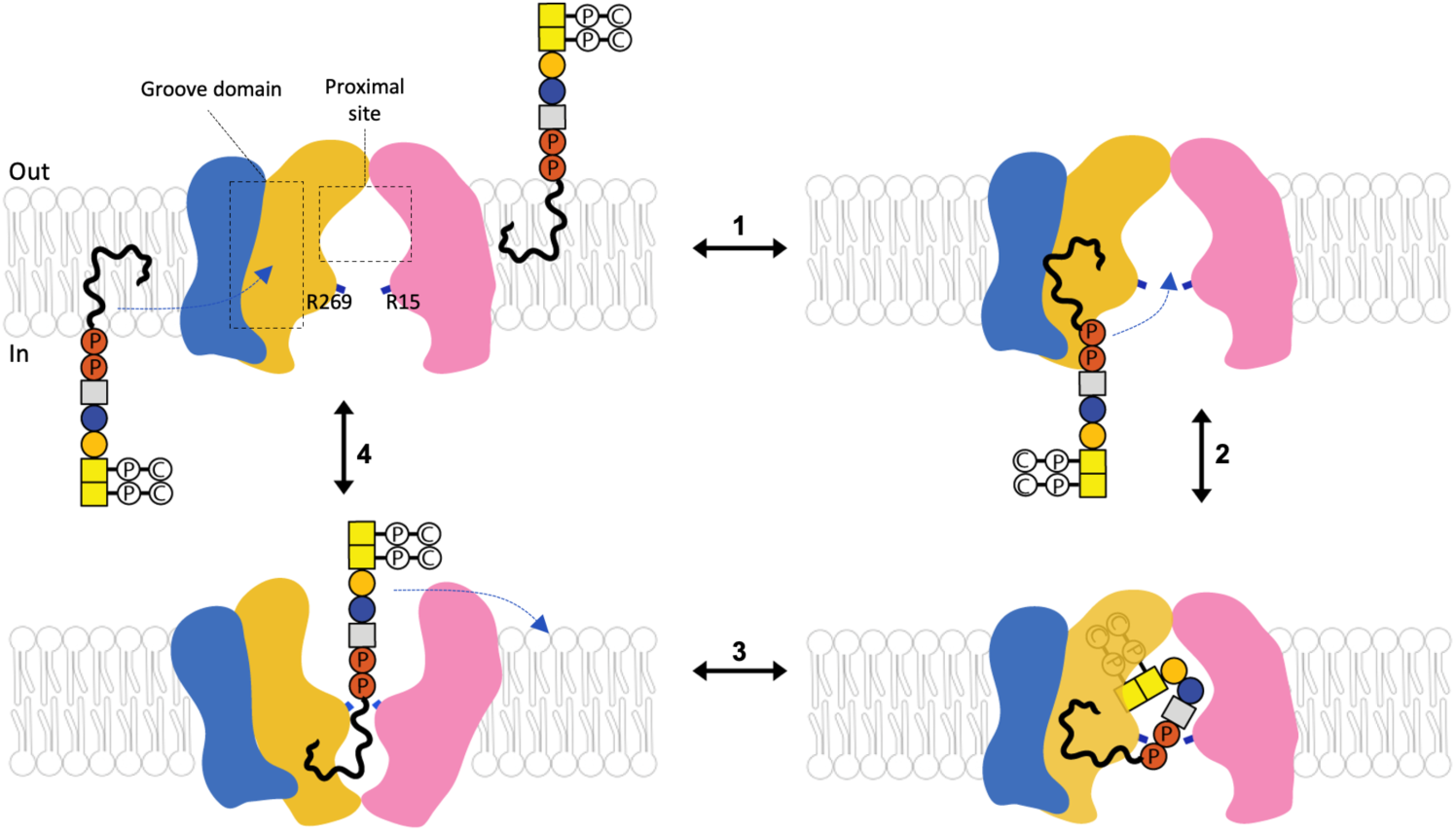
Mechanistic model of teichoic acid flipping by TacF. The undecaprenyl tail of the teichoic acid is recognized through hydrophobic interactions at the groove domain formed by TM13 and TM14 (**1**), promoting binding of the diphosphate group by R15 and R269 located at the TM helices forming the lateral entrance (TM1 and TM8) (**2**). The teichoic acid repeating unit enters the central cavity and is coordinated by multiple interactions with residues in the proximal site (**2**). TacF transitions to the outward-facing state, facilitating the release of the teichoic acid into the extracellular leaflet of the membrane (**3**). TacF resets to the inward-facing state before initiating a new cycle (**4**). N– and C-terminal domains are colored pink and yellow, respectively. TM13 and 14 are colored blue.

The recognition mode of teichoic acid by TacF resembles the proposed model for how the flippase MurJ recognizes its lipid-II substrate^51^, for which it has been shown that the groove domain and positively charged residues positioned in TM1 and TM8 stabilize the binding of the undecaprenyl-diphosphate moiety of lipid-II, while the proximal and the distal sites contribute to recognizing the GlcNAc-MurNAc and pentapeptide moieties, respectively^51^. In the case of Wzx, it has been suggested that it binds the undecaprenyl tail of its polyprenyl-diphosphate oligosaccharide substrate through interactions with a hydrophobic groove outside the central cavity^77^. However, this groove is distinct from that observed in TacF and MurJ due to the absence of TM13 and 14 ^77^. Similar to TacF and MurJ, Wzx recognizes the diphosphate group of its substrate through positively charged residues located at the entrance of the transport channel, while the oligosaccharide headgroup binds inside the central cavity^77^. The mechanistic analysis of TacF presented here, together with the prior work on MurJ^48,49,80,81^ and Wzx^77^, supports a conserved substrate recognition mechanism among MOP flippases, but studies on additional members of this superfamily are required to establish a general model of transport by flippases in this superfamily.

Previous studies have shown that strains carrying specific mutations win the *tacF* gene exhibit promiscuous activity of TacF towards unmodified teichoic acids, allowing growth of *S. pneumoniae* in the absence of choline^23,40,42,83^. The mutations reported involved residues V32, I100, P104, F106, F107, F209, A214, V234, I247, F296, and I298, with a minimum of two residue changes required to allow growth of *S. pneumoniae* in choline-free medium^42^ (**Suppl. Fig. 8B**). Interestingly, none of these residues are part of the domains critical for recognizing the teichoic acid molecule. Four of these residues are located at the interface between the N– and C-terminal domains on the extracellular side of TacF (V32, I247, V234, F107) (**Suppl. Fig. 8B**), so we suggest that their effect may come from affecting the rocker-switch transition from inward– to outward-facing conformations, but further studies are needed to confirm this. On the other hand, two residues located in the groove domain (F296 and I298) (**Suppl. Fig. 8B**) may influence substrate selectivity by directly affecting teichoic acid recognition.

Our findings show that the hydrophobic properties of the groove domain are broadly conserved among TacF homologues, while variations in the proximal and distal sites may reflect specific adaptations to different teichoic acid substrates. Multiple *Streptococcus* species in cluster I (**Figure 2**) carry canonical phosphocholine-modified teichoic acid backbones^23,58,59^, unlike species in clusters II to V, where this modification has not been detected^2,60–62,84^. This suggests that TacF proteins in clusters II to V may specialize in recognizing teichoic acid backbones through different mechanisms than the one described here for *S. pneumoniae* TacF.

The flipping of teichoic acid represents a rate-limiting step in the cell wall biosynthesis pathway in *S. pneumoniae*, making it a promising therapeutic target. Our integrated characterization of TacF provides fundamental insights into the molecular mechanism by which it transports phosphocholine-modified teichoic acids and advances our understanding of the flipping mechanism of MOP superfamily flippases. This study may contribute to the development of novel therapies targeting multi-resistant *S. pneumoniae* strains.

## Materials and Methods

### Preparation of BRIL-TacF constructs

A synthetic gene fragment containing the *Streptococcus pneumoniae tacF* gene was cloned into a modified pET-19b vector (Novagen) yielding the wild-type TacF construct. This incorporated an N-terminal polyhistidine tag flanked by a TEV cleavage site. The same vector was utilized for the design of the BRIL-TacF fusion partners. The first 12 N-terminal residues of TacF were truncated to allow BRIL insertion, immediately preceding the first transmembrane helix. A total of six BRIL-TacF constructs were generated: BRIL_5_-TacF, BRIL_7_-TacF, BRIL_8_-TacF, BRIL_9_-TacF, BRIL_10_-TacF, and BRIL_12_-TacF, where 5, 7, 8, 9, 10, and 12 residues from the N-terminal domain of TacF were removed, respectively (**Suppl. Table 1**).

### Protein expression and purification of TacF WT and BRIL-TacF constructs

Competent *E. coli* BL21 (DE3) cells were transformed and grown at 37°C in Terrific Broth (TB) media supplemented with 0.4% glycerol. Protein expression was induced with 0.4mM isopropyl β-D-1-thiogalactopyranoside (IPTG) followed by incubation at 37°C for 1 hour. Cell pellets were harvested and stored at –80°C. For membrane isolation, pellets were resuspended by continuous stirring at 4°C for 2h in lysis buffer (50 mM Tris-HCl, pH 8.0, 200 mM NaCl, 3 mM β-mercaptoethanol, and 0.5 mM PMSF). Cell membranes were isolated by differential centrifugation and resuspended in resuspension buffer containing 50 mM Tris-HCl, pH 8.0, 200 mM NaCl, and 3 mM β-mercaptoethanol. For purification, membranes were solubilized in buffer containing 50 mM MES pH 6.5, 200 mM NaCl, 20 mM Imidazole, 15% Glycerol, 2 mM β – mercaptoethanol, and 1% DDM, prior to centrifugation and loading into a pre-equilibrated nickel-nitrilotriacetic acid (Ni-NTA) column, pre-equilibrated with 50 mM MES pH 6.5, 150 mM NaCl, 20 mM Imidazole, 10 % glycerol, 3 mM β –mercaptoethanol, and 0.02% DDM. The column was washed with buffer containing 50 mM MES pH 6.5, 150 mM NaCl, 50 mM Imidazole, 10 % glycerol, 3 mM β-mercaptoethanol, and 0.02% DDM. The protein was eluted with buffer containing 50 mM MES pH 6.5, 150 mM NaCl, 400 mM Imidazole, 10 % glycerol, 3 mM β – mercaptoethanol, and 0.02% DDM. The eluted protein buffer was exchanged using PD-10 desalting columns (GE Healthcare) to 20 mM MES pH 6.5, 150 mM NaCl, and 0.02% DDM prior to incubation with Tobacco Etch Virus (TEV) protease. After TEV removal, the protein was concentrated using 50kDa cutoff Vivaspin concentrators and injected onto a Superdex 200 Increase 10/300 column previously equilibrated with buffer containing 20 mM MES pH 6.5, 150 mM NaCl, and 0.02% DDM in a ÄKTA Pure system (GE Healthcare).

### Thermostability assays

Purified BRIL_i_-TacF proteins at 0.2mg/mL were incubated at different temperatures (4 °C, 22 °C, 42 °C, 65 °C, and 90 °C) for 15 minutes. Following the heat treatment, samples were centrifuged at 13,300 g for 15 minutes at 4°C. The proteins were transferred to a 96-well plate and injected into a Superose 6 10/300 GL column (GE Healthcare) pre-equilibrated with buffer containing 20 mM MES pH 6.5, 150 mM NaCl, and 0.02% DDM. Chromatograms were analyzed, and the main peak height was plotted against the incubation temperatures using GraphPad Prism 8.

### Mass photometry analysis

Mass photometry experiments of nanodisc-reconstituted TacF constructs and empty nanodiscs were carried out using a Refeyn TwoMP mass photometer^85^. Recordings were carried out at 20nM protein concentration and in buffer containing 50 mM Tris-HCl pH 8.0, 150 mM NaCl. Each sample was loaded onto a pre-focused buffer droplet placed on a cover slide (Marienfeld) secured by a silicone gasket (Grace Biolabs). Interference signals were recorded for 1 minute immediately after mixing. The mass photometry data were initially recorded in ratiometric contrast units and subsequently converted to molecular mass units using a calibration curve generated from standard proteins of known molecular weights (Bovine Serum Albumin –66 kDa-and Bovine Thyroglobulin –330 kDa monomer, 660 kDa dimer-). The acquired movies were processed using the manufacturer’s software (DiscoverMP) and plotted using GraphPad Prism 8.

### Reconstitution of TacF in nanodiscs

Purified TacF variants were reconstituted into MSP1D1 nanodiscs composed of a 3:1 molar ratio of POPG to DAG (Avanti Lipids). Reconstitution was performed at a TacF:lipid:MSP1D1 molar ratio of 1:60:3 using a reconstitution buffer composed of 50mM Tris-HCL pH 8.0, 50mM NaCl, and 10% glycerol. Detergent was removed by incubation with Bio-Beads (SM2, Bio-Rad) for 16 hours under gentle rotation at 4°C. After incubation, the sample was centrifuged at 13,000 g for 5 minutes at 4°C and the supernatant was loaded into a Superdex 200 Increase 10/300 column equilibrated with buffer containing 50 mM Tris-HCl pH 8.0, and 150 mM NaCl. The fractions corresponding to the peak of TacF reconstituted in nanodiscs were collected and used for Cryo-EM experiments.

### Purification of BAG2 antibody fragment (Fab)

The expression and purification of BAG2 was carried out as previously described^86^, with slight modifications. BAG2 was recombinantly expressed in *E. coli* BL21 (DE3) cultured at 37°C in 2x Yeast extract Tryptone medium (2YT) media. Protein expression was induced with 1mM IPTG upon reaching an O.D._600_ of 0.6 and further incubated for 5 hours. For protein purification, cell pellets were resuspended in lysis buffer containing 20 mM Tris-HCl pH 7.5, 150 mM NaCl, 1 mM PMSF, and 10 mg/L DNAse I. Resuspended cells were disrupted using a high-pressure homogenizer and the resulting lysate was subjected to heat precipitation at 60°C for 30 minutes, followed by centrifugation at 50,000 g for 30 minutes at 4°C. The supernatant was later filtered through a 0.22μm membrane filter and injected into a Protein L column (GE Healthcare) pre-equilibrated with running buffer containing Tris-HCl pH 7.5, and 500 mM NaCl, using an ÄKTA Pure system. The column was washed with 10 column volumes of running buffer. BAG2 was eluted with 0.1M acetic acid in 2mL fractions. Eluted BAG2 was loaded onto a Resource S column (GE Healthcare) pre-equilibrated with buffer A (50 mM sodium acetate pH 5.0) and subjected to cation-exchange chromatography. Unspecific interactions were washed away with buffer A and the protein was eluted using a salt gradient (0-100%) in buffer composed of 50 mM sodium acetate pH 5.0, and 2M NaCl. Eluted protein was then desalted into a buffer containing 50 mM Tris-HCl pH 7.5, 150 mM NaCl, aliquoted, flash frozen and stored at –80°C.

### Cryo-EM data collection and processing

Nanodiscs reconstituted BRIL_9_-TacF:BAG2 complex were subjected to size exclusion chromatography in a Superdex 200 Increase 10/300 column. The peak fraction was further concentrated to 0.8-1mg/mL and applied to glow-discharged copper holey carbon R1.2/1.3 300-mesh grids (Quantifoil). Freezing conditions were set to 95% humidity and 4°C. Grids were blotted before vitrification by plunging in liquid ethane using a Mark IV Vitrobot (Thermo Fisher Scientific). Cryo-EM data was collected using a Glacios Cryo 200kV TEM (Thermo Fisher Scientific) equipped with a K3 direct electron detector (Gatan) and operated using serialEM^87^. Micrographs were collected with a defocus range of 0.5 to 3 μm and at 0.878 Å/pixel at a nominal magnification of 46,000x. The total electron dose was 55 e−/Å^2^. The overall data processing workflow is illustrated in **Supplementary Fig. 3**. Data processing was carried out entirely in cryoSPARC v4.7.1 ^88^. Beam-induced drift correction was applied to 6,591 raw movies following patch motion correction. The micrographs were then manually curated and a small subset of 10 micrographs was used to optimize particle-picking parameters through blob-picking, selecting particles between 100 and 200Å in diameter. Particles were later extracted from the entire data set, reducing the box size by Fourier cropping. After multiple iterations of 2D classification, the selected particles were used to generate ab initio reconstructions. Particles corresponding to the best reconstructions were subjected to multiple iterations of heterogeneous refinement using the best ab initio reconstruction as a target and a low-quality maps as a decoy. The generated electron density maps were further refined with nonuniform (NU) refinement jobs. The highest resolution NU refinement map was used to generate a template for automated template-based particle picking. Picked particles were processed using the same protocol and subjected to multiple rounds of 3D-calssificatoon, followed by masking of the nanodisc and part of the BAG2 Fab. This, together with local refinement helped to increase the resolution, resulting in a 3.63Å local resolution map. Model building was carried out using Coot^89^. Figures of models and maps were made using PyMOL (The PyMOL, Molecular Graphics Systems, Schödinger LLC) and ChimeraX^90^.

### MD simulations

TacF was simulated in a heterogeneous bilayer composed of POPG (40%), cardiolipin (CL) (40%), and 1-palmitoyl-2-oleoyl-3-O-(β-d-glucosyl)-sn-glycerol (BGLC-DAG-PO) (20%) using the modified scripts from the CHARMM-GUI web server^91^. Sodium and chloride ions were added to a total ionic concentration of 150mM. An all-atom CHARMM36m force field was used for proteins, lipids and ions, and the TIP3P model for water molecules^92,93^. The force field parameters for the teichoic acid molecule were derived by modifying pre-parameters established for a structurally related lipid-linked oligosaccharide^94^. MD trajectories were analyzed using MDAnalysis and in-house scripts^95^. All simulations were performed using GROMACS 2024^96,97^. The initial setups were energy-minimized for 5,000 steepest descent steps and equilibrated for 1.5ns in a canonical (NVT) ensemble, followed by 7ns in an isothermal-isobaric (NPT) ensemble under periodic boundary conditions. Restraints on the positions of non-hydrogen protein atoms of initially 4,000 kJ·mol^−1^·nm^2^ were gradually released during equilibration. The cutoff distance for non-bonded interactions was set to 1.2nm. Particle-mesh Ewald summation was employed to handle long-range electrostatic interactions^98^, using cubic interpolation and a grid spacing of 0.12nm. The time step was initially set to 1fs during the NVT equilibration and increased to 2fs during the NPT equilibration. The LINCS algorithm was used to fix bond lengths^99,100^. During the equilibration phase, constant temperature and pressure were established with a Berendsen thermostat combined with a coupling constant of 1.0ps and a semi-isotropic Berndsen barostat with a compressibility of 4.5 ×10^−5^ bar^−1 101^. The Berendsen thermostat and barostat were replaced by a V-rescale thermostat^102^ and a C-rescale barostat^103^ during production runs.

### Direct Coupling Analysis

A direct coupling analysis (DCA) of the TacF protein sequence was performed using the EVCouplings software package^73^. The full-length amino acid sequence of TacF (UniProt code Q8DPI1) and a multiple sequence alignment comprising 138,448 sequences generated using JackHMMER^104^, was submitted as a query to the EVCouplings webserver to generate DCA coupling scores (CN scores) with probability scores. Out of a total of 105,112 residue pairs obtained, the top 651 scored pairs were considered long range, with constituent residues that were greater than five positions apart in the amino acid sequence, and having a probability of 0.99 or greater. The top 100 of these long-range pairs, ranked by CN score, were analyzed and visualized on the TacF structure using PyMOL software.

### SSN analysis

To perform sequence similarity networks (SSNs) analysis of TacF homologues, protein sequences of interest were submitted to the EFI-EST webserver^105^, where pairwise sequence similarities were calculated using BLAST and transformed into SSN files. After selecting percent alignment thresholds to control the network resolution, the resulting XGMML file was imported into Cytoscape^106^. The SSNs were represented in Cytoscape according to sequence similarity or according to the bacterial species where the sequence is found, with each node representing a protein sequence or a collection of sequences with more than 95% identity, whereas edges denote sequence similarity above a 40% identity cutoff.

### *S. pneumoniae* growth strains

All strains used in this study were derived from the clinical isolated serotype 2 *S. pneumoniae* D39V^107^. C+Y media at pH 6.8, adapted from Adams and Roe^108^, was used as growth medium. Genomic DNA Golden Gate assembly plasmids were used to transform *S. pneumoniae* after induction with competence stimulation peptide 1 (CSP-1) as previously described^108^. Transformants were selected on Columbia agar with 2% sheep blood at 37°C and 5% CO_2_ as well as an antibiotic mix (4.54 μg/ml chloramphenicol, 0.5 μg/ml erythromycin, 250 μg/ml kanamycin, 100 μg/ml spectinomycin). Strains were sequenced by sanger sequencing at Microsynth and stocked at O.D._595_ 0.3 at –80°C with 14.5% glycerol.

### Microplate growth assay

To perform growth assays, the bacterial strains were cultured in fresh C+Y media at pH = 6.8 with or without 1 mM IPTG and/or 50 ng/mL aTc at 37°C 5% CO_2_ until reaching O.D._600_ 0.2. The cultures were then diluted to O.D._600_ 0.01 in fresh C+Y medium (pH = 6.8) with or without 1 mM IPTG and/or 50 ng/mL aTc, and dispensed into a 96-well plate (200 µl per well). Cell growth was monitored by measuring optical density at 595nm every 10 minutes for 16 hours using a microplate reader (TECAN infinite F200 Pro). Each growth assay was performed in triplicate, and the mean value was plotted, with the SEM (Standard Error of the Mean) represented by an area around the curve. Each well was normalized by subtracting the minimum value of each well over the period. To generate the barplots, the area under the curve was calculated from 0h to 8h, in an empirical manner, where O.D._600_ values are summed over the time. Plots were made using BactEXTRACT (https://doi.org/10.1099/acmi.0.000742.v3).

## Data Availability

The data that support the findings of this study are available from the corresponding author upon reasonable request.

## Acknowledgement

We thank Professor Anthony Kossiakoff and Dr. Somnath Mukherjee from the University of Chicago for the generous gift of BRIL and BAG2 constructs and protocols. We thank the staff at the electron microscopy facility at Biozentrum of the University of Basel (BioEM lab) for facilitating data collection. Work in the Perez lab is supported by the Swiss National Science Foundation (SNSF) (310030_207974) and NIH grant R35GM158187. Work in the Veening lab is supported by SNSF grants 310030_200792 and 51NF40_180541. The Mehdipour lab acknowledges funding from the Special Research Fund of Ghent University, grant number BOF.STG.2021.0037.01. The Avci lab acknowledges funding from the NIH (R01AI123383, R01AI152766). NK acknowledges funding from NIH/NIGMS (R35-GM139656).

## Author Contributions

G.C. designed constructs, established purification conditions of TacF-WT and BRIL-TacF, expressed and purified proteins, and prepared the sample for cryo-EM studies. A.C. expressed and purified proteins for characterization by mass photometry of BAG2/BRIL-TacF interaction, and performed thermostability assays of BRIL-TacF constructs. G.C. performed characterization by mass photometry of TacF-WT. G.C. and C.P. prepared grids for cryo-EM data collection and performed preliminary cryo-EM data processing and model building. E.C. and C.P. performed cryo-EM data processing, structure building, and analysis of the deposited structure. E.C. and C.P. performed and analyzed SSN data. J.D. performed complementation and growth assays under supervision from J.W.V. A.T.M. performed MD simulations under supervision of A.R.M. M.C. performed chemical synthesis of lipids under supervision of J.L.R. R.B. and A.C. performed evolutionary analysis under supervision of N.K. and C.P. E.S.D. performed complementation and growth assays under supervision of F.Y.A. A.C. and C.P. wrote the manuscript. C.P. conceived the project.

## Author Information

Competing interests: J.W.V. is a scientific advisory board member at i-Seq Biotechnology. The remaining authors declare no competing interests.

Data and materials availability: Electron microscopy density maps and atomic models have been deposited in the EMDB and PDB, respectively, with accession codes EMD-72367 and PDB 9XZN. Molecular dynamics simulations performed in this study have been deposited in Zenodo: doi 10.5281/zenodo.17151412.

