## Supplementary Information for "Mechanistic basis of teichoic acid transport by a gatekeeper flippase"

**This PDF file includes:**

Figs. S1 to S9

Table S1

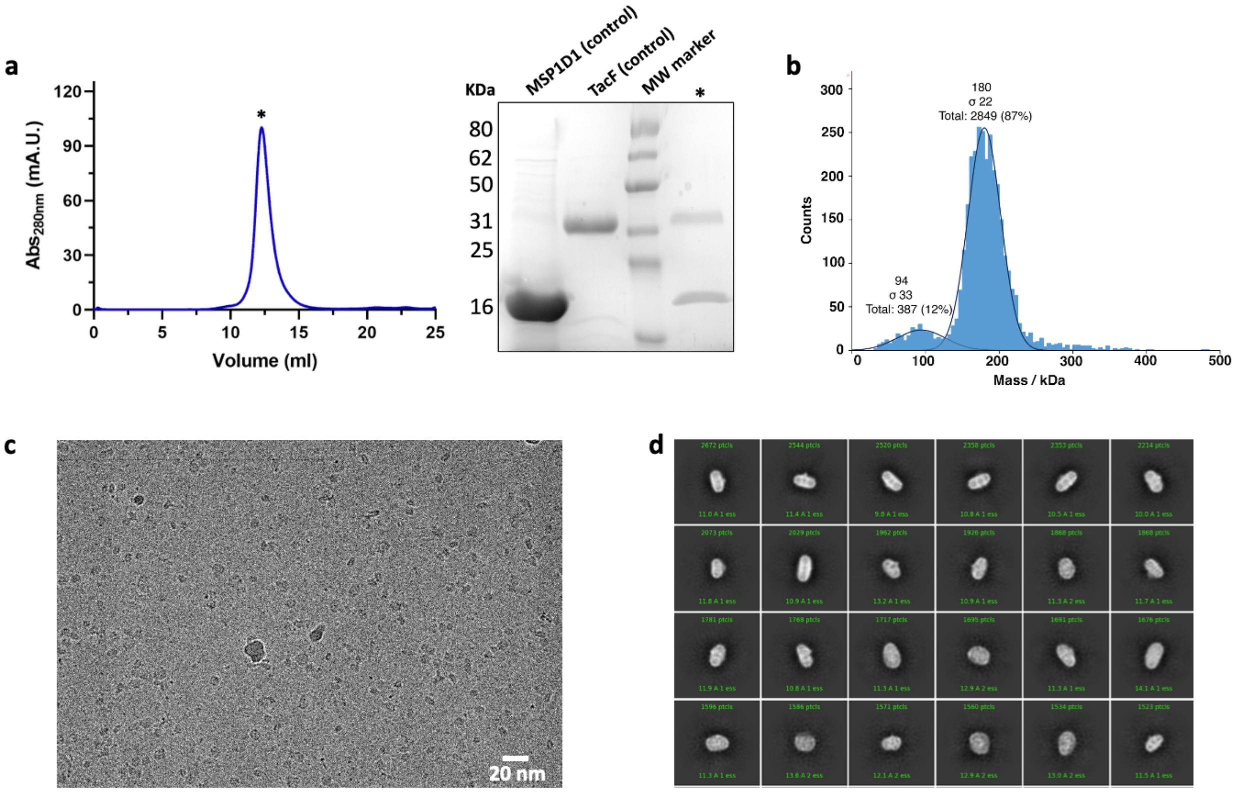


**Supplementary Figure 1**. **Purification and cryo-EM analysis of TacF wild-type**. **A**. Size exclusion chromatography profile in a Superdex 200 Increase 10/300 column for TacF reconstituted in MSP1D1 nanodiscs. SDS-PAGE of the main peak and controls (purified MSP1D1 and TacF) is shown. **B**. Mass photometry analysis of TacF reconstituted in MSP1D1 nanodiscs. The two distinct populations indicate empty nanodiscs (94 +/- 33kDa) and TacF reconstituted nanodiscs 180 +/- 22kDa). **C**. Representative micrograph of TacF nanodiscs. **D**. 2D class averages of TacF nanodiscs.

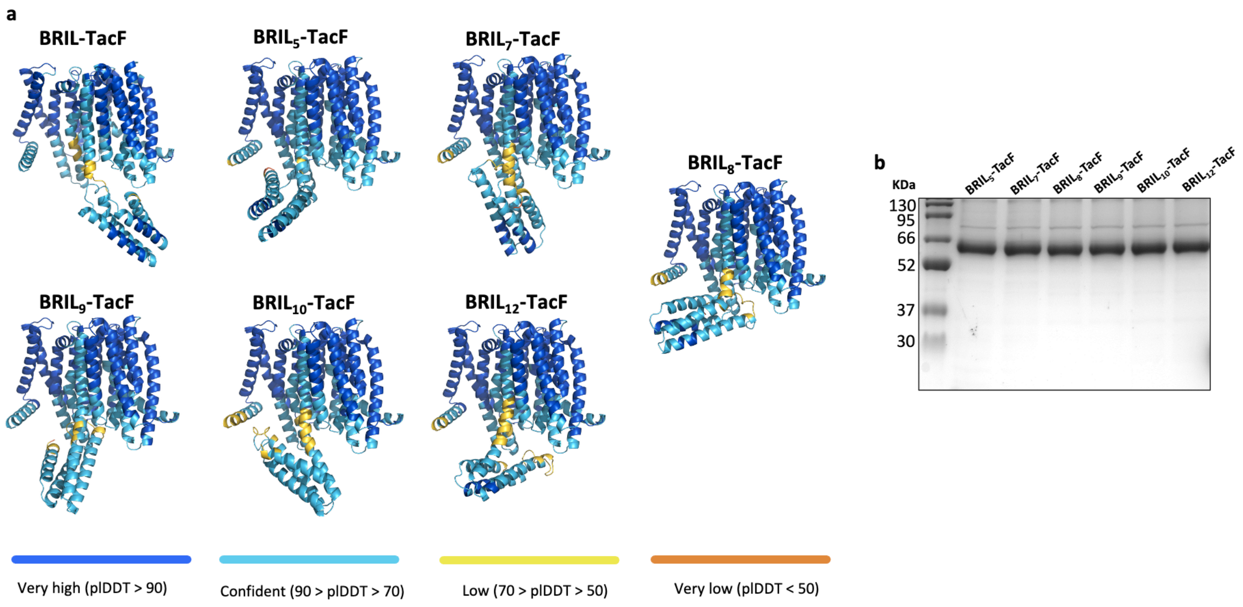


**Supplementary Figure 2. *In silico* analysis of BRIL-TacF constructs**. **A**. AlphaFold-3 predicted models of BRIL–TacF fusion constructs. The predicted local distance difference test (pLDDT) indicates local confidence per residue. **B**. SDS-PAGE analysis of purified BRIL-TacF constructs.

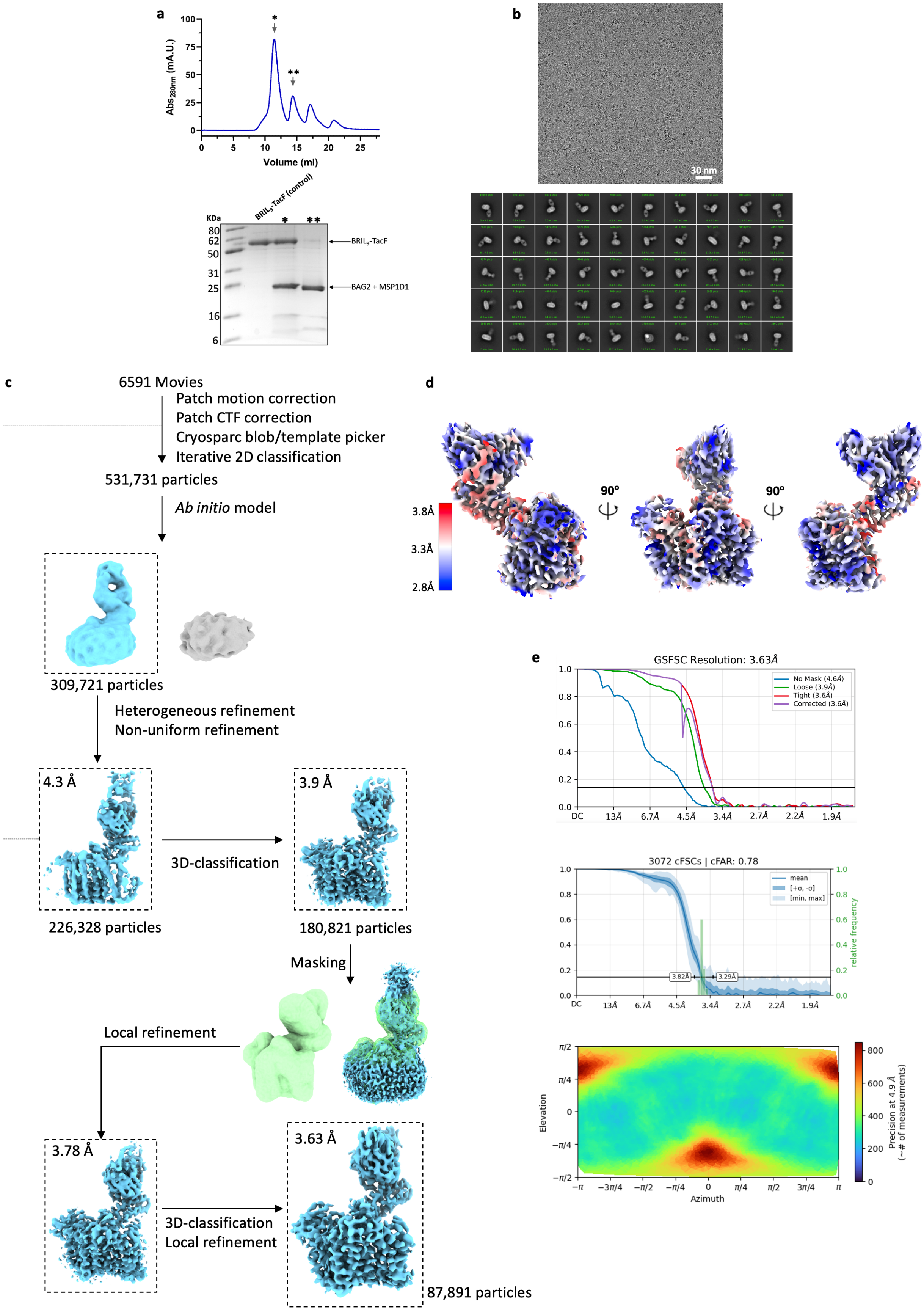
a
Abs280nm (mA.U.)
Volume (ml)
BRIL5-TacF (control)
KDa
BRIL5-TacF
BAG2 + MSP1D1
b
30 nm
c
6591 Movies
Patch motion correction
Patch CTF correction
Cryosparc blob/template picker
Iterative 2D classification
531,731 particles
Ab initio model
309,721 particles
Heterogeneous refinement
Non-uniform refinement
4.3 Å
226,328 particles
3D-classification
3.9 Å
180,821 particles
Masking
Local refinement
3.78 Å
3D-classification
Local refinement
3.63 Å
87,891 particles
d
3.8Å
3.3Å
2.8Å
90°
90°
e
GSFSC Resolution: 3.63Å
No Mask (4.6Å)
Loose (3.9Å)
Tight (3.6Å)
Corrected (3.6Å)
3072 cFSCs | cFAR: 0.78
mean
[+σ, -σ]
[min, max]
relative frequency
3.82Å
3.29Å
Elevation
Azimuth
Precision at 4.9 Å (~# of measurements)

**Supplementary Figure 3. Cryo-EM characterization of BRIL$_9$-TacF in lipid nanodiscs**. **A**. Size exclusion chromatography profile in a Superdex 200 Increase 10/300 column for BRIL$_9$-TacF:BAG2 complex reconstituted in MSP1D1 nanodiscs. SDS-PAGE of the main peaks are shown. **B**. Representative micrograph of BRIL9-TacF:BAG2 nanodiscs and 2D class averages. **C**. Cryo-EM data processing workflow. A set of 6,591 movies was processed, resulting in a set of 531,731 particles that was used as input to generate an *ab initio* reconstruction volume, which was refined through heterogeneous and non-uniform refinement. The resulting volume was used to pick a new set of particles, which were further processed using 3D classification, local refinements, and masking of the nanodisc and part of the BAG2 Fab. This resulted in a 3.63 Å local resolution map. **D**. Local resolution map of the final volume computed in cryoSPARC. **E**. GSFSC and cFSC curves used for resolution estimation of the final map. *Bottom*, Angular viewing directions distribution of particles contributing to the final map.

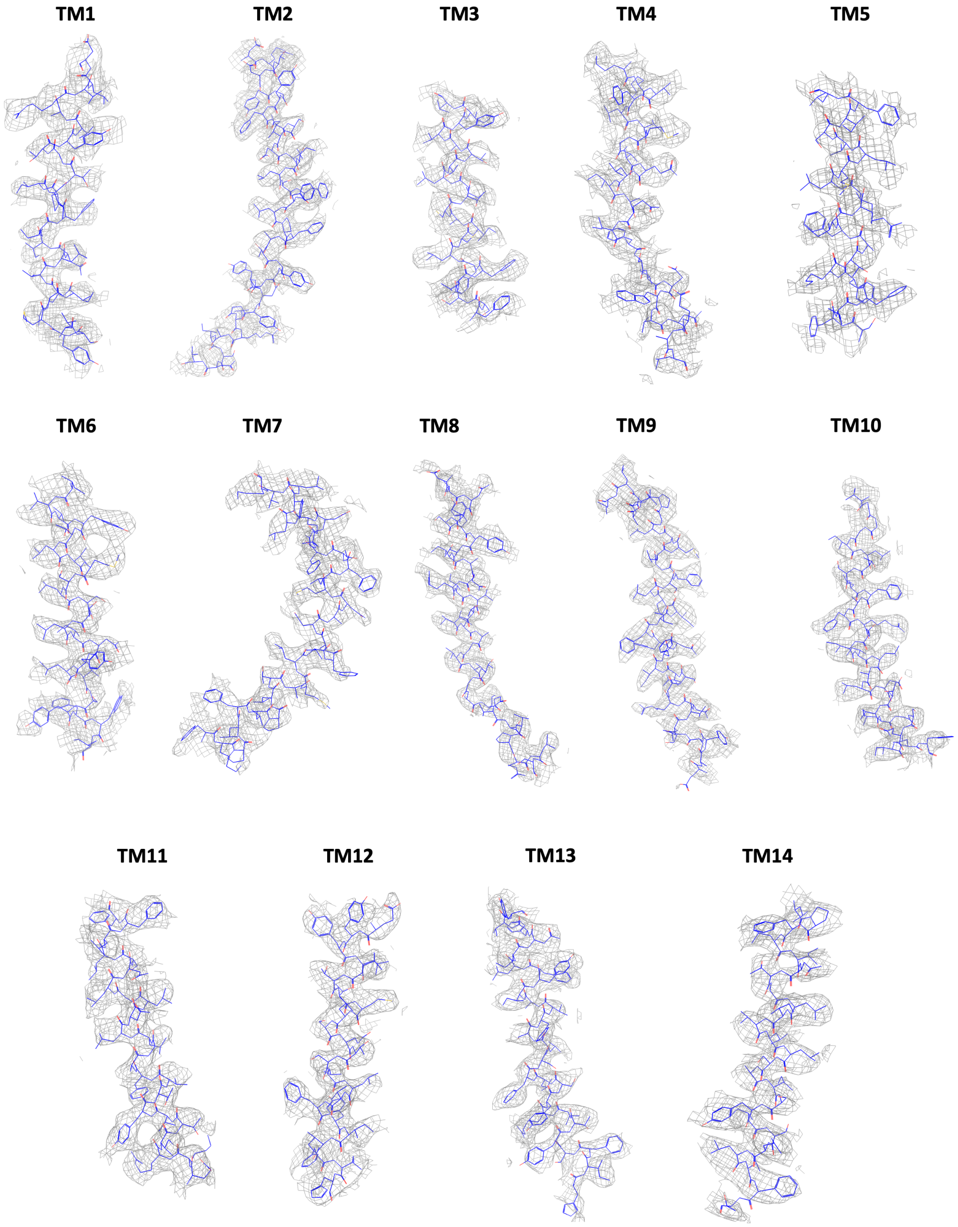


**Supplementary Figure 4. Cryo-EM density of the refined TacF structure**. The TacF model of TM1 through 14 is shown using line representation. Cryo-EM volume density is represented as a grey mesh.

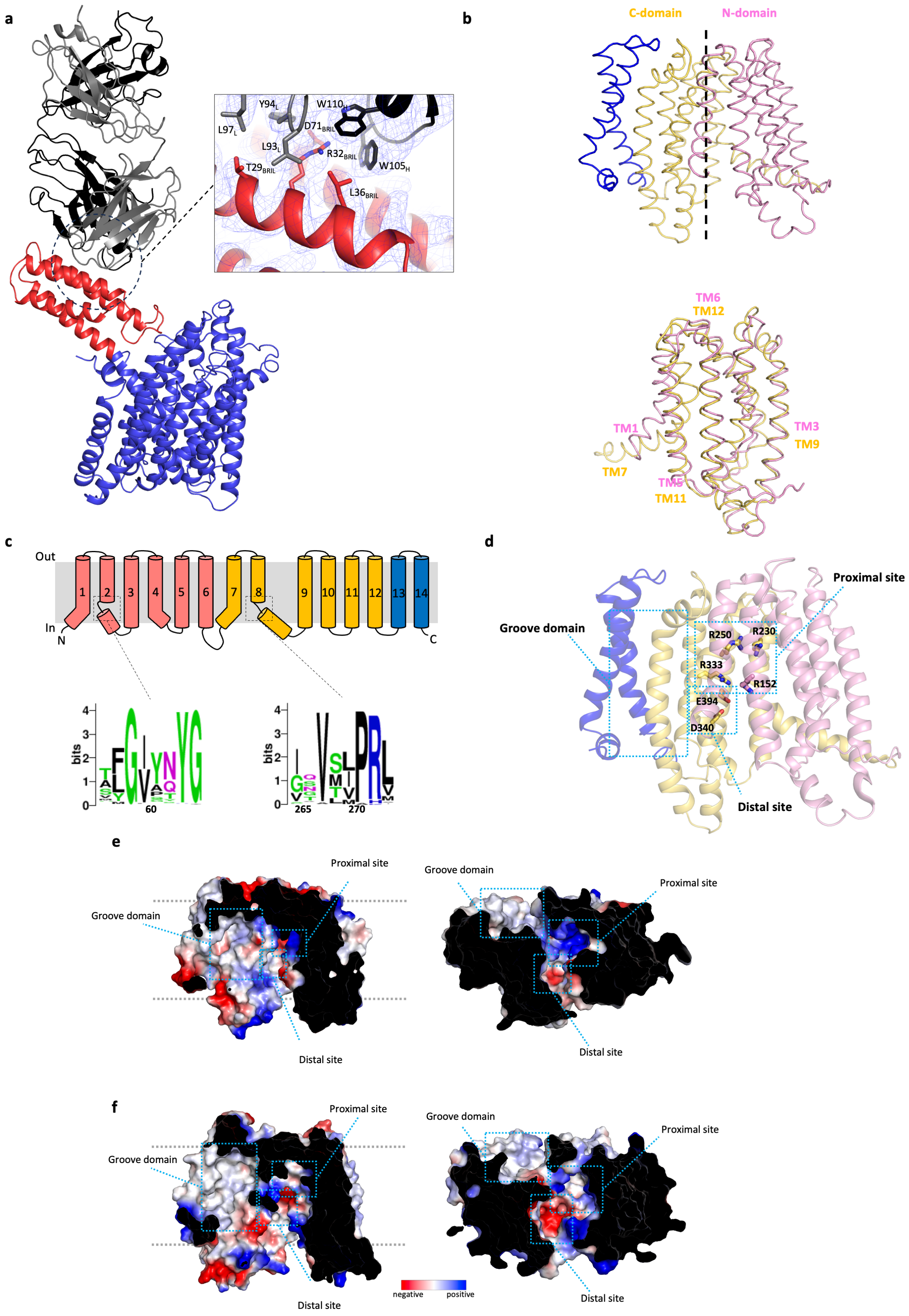
a
Y94L
W110H
L97L
D71BRIL
L93L
R32BRIL
T29BRIL
W105H
L36BRIL
b
C-domain
N-domain
TM6
TM12
TM1
TM7
TM3
TM9
TM5
TM11
c
Out
In
N
C
1
2
3
4
5
6
7
8
9
10
11
12
13
14
bits
60
265
270
d
Proximal site
Groove domain
R250
R230
R333
R152
E394
D340
Distal site
e
Proximal site
Groove domain
Distal site
f
Proximal site
Groove domain
Distal site
negative
positive

**Supplementary Figure 5. Analysis of the TacF structure and topology**. **A**. The structure of $BRIL_9$-TacF (red and blue, respectively) in complex with BAG2 Fab (Grey) is shown. *Inlet* shows the interactions involved in the binding of BAG2 to the BRIL fragment. The cryo-EM map is displayed as a blue mesh. **B**. Pseudosymmetry of the N- and C-terminal domains. *Bottom*, N-terminal (pink ribbons) and C-terminal (yellow ribbons) are superimposed. **C**. Topology of TacF based on the cryo-EM structure and sequence logos analysis of unwound segments observed in TM2 and TM8. **D**. Residues contributing to the positive surface of the proximal site and the negative charge of the distal site. **E** and **F**. Side and cytoplasmic views of a surface electrostatic potential representation of MurJ (PDB ID: 5T77) (**E**) and an AlphaFold model of Rft1 (**F**), showing the locations of the proximal and distal sites, as well as the groove domain.

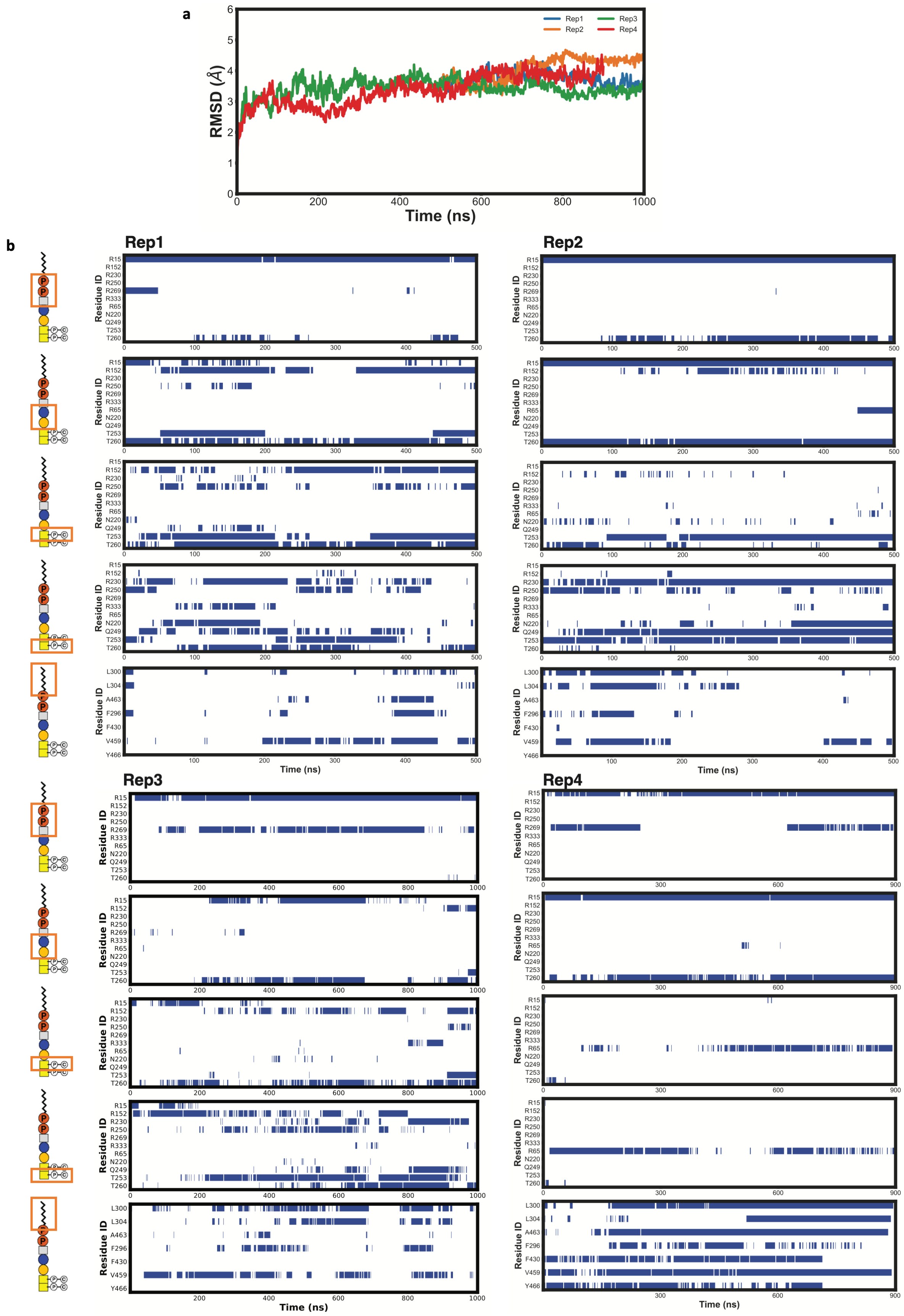

a
RMSD (Å)
Time (ns)
Rep1
Rep2
Rep3
Rep4
b
Residue ID
Time (ns)

**Supplementary Figure 6. MD simulations of TacF and interactions with teichoic acid**. **A**. The root-mean-square-deviation (RMSD) of TacF starting from the equilibration structure during the different simulation replicates. **B**. The time evolution of the interaction of teichoic acid with residues in the binding cavity. Interactions between teichoic acid and an amino acid were considered if at least one pair of their non-hydrogen atoms was within 4 Å of each other. The interactions are shown for the five different parts of the teichoic acid separately (chemical structure representation of teichoic acid as shown in **Fig. 1A**).

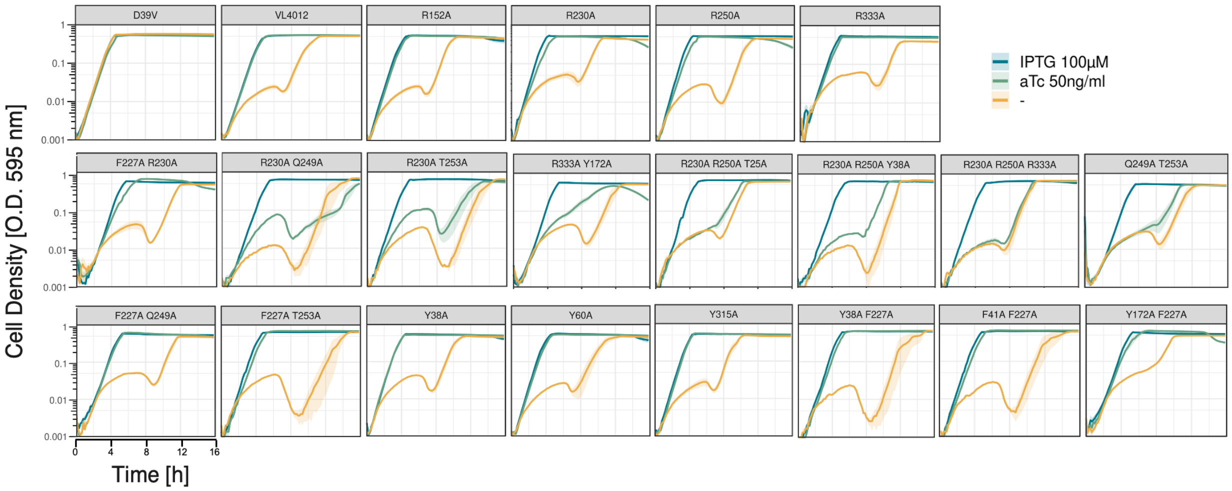


**Supplementary Figure 7. Growth curves of *S. pneumoniae* strains carrying variants of TacF**. 16h growth curves of *S. pneumoniae* D39V strains carrying the double expression system for the *tacF* wild type allele (VL4012) or *tacF* variants are shown. Cultures were grown in media supplemented with IPTG (blue), aTc (green), or no inducer (yellow). *S. pneumoniae* D39V growth curve indicates growth of the unmodified D39V strain.

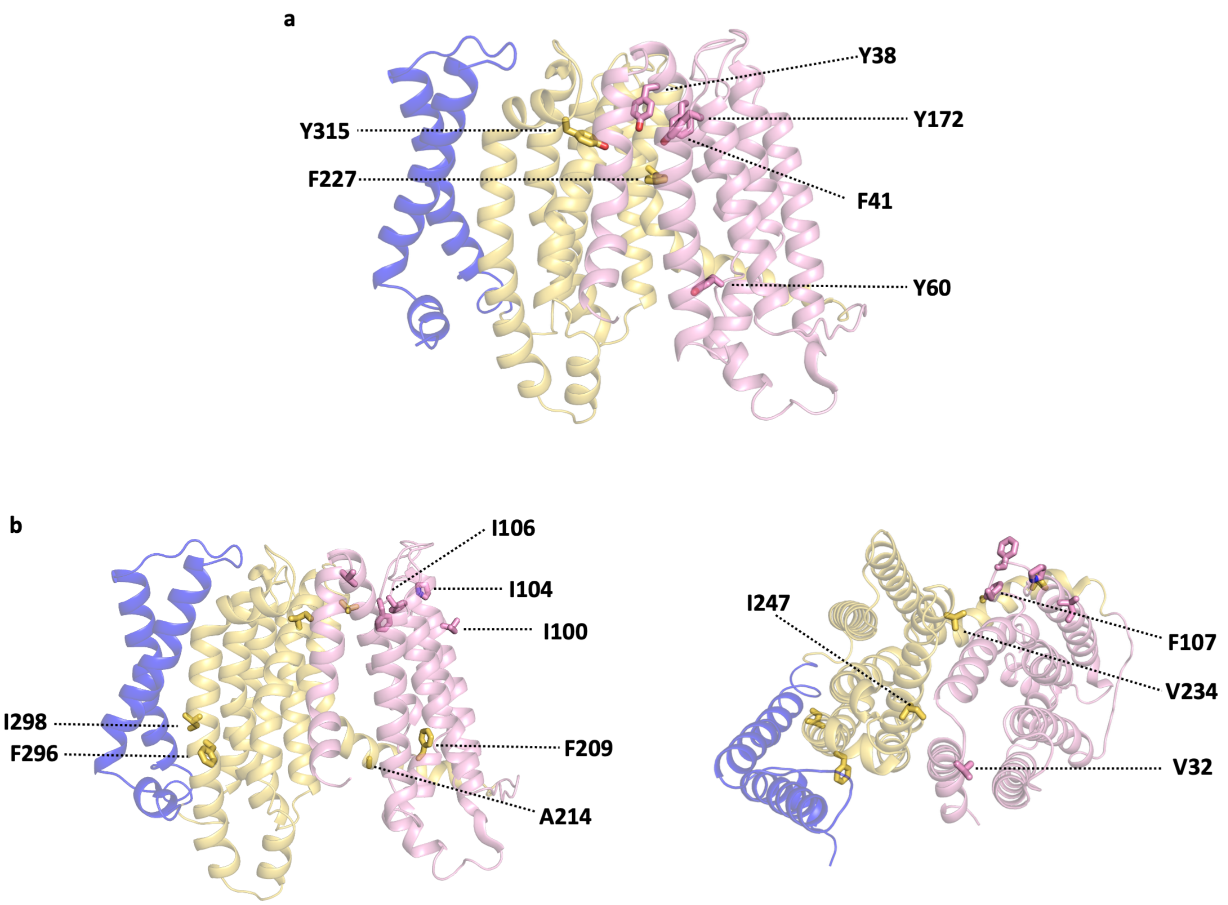


**Supplementary Figure 8. Analysis of the TacF structure**. **A**. Aromatic residues located in the central cavity of TacF. **B**. Residues that, when mutated, have been previously reported to induce promiscuous activity of TacF towards teichoic acids lacking phosphocholine[23,40,42,83].

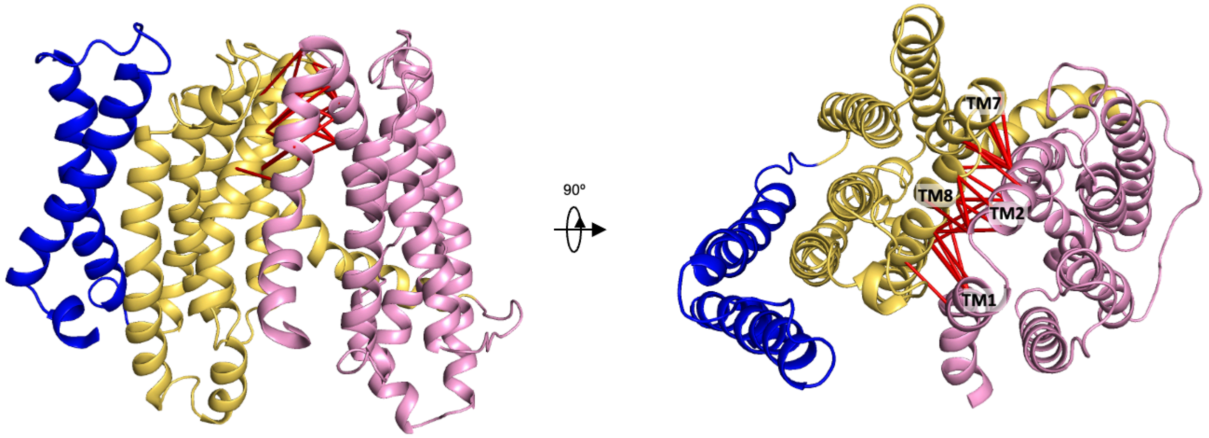


**Supplementary Figure 9. Evolutionary coupling analysis of TacF**. Coevolving pairs (red lines) with short inter-residue distances on the extracellular side of the cryo-EM TacF structure.

**Supplementary Table 1. Primers used to generate the BRIL-TacF constructs**

| Primer | Sequence | $T_m$ (°C) | Reference |
|---|---|---|---|
| BRIL$_5$-TacF For | 5'- CAT ATA TCC AGA AGT ATC TTA AAC TGA ACG CCC TGA GCT ATA TGG GTA TTC-3’ | 68.7 | This study |
| BRIL$_7$-TacF For | 5'- CAT ATA TCC AGA AGT ATC TTA ACG CCC TGA GCT ATA TGG GTA TTC-3’ | 66.9 | This study |
| BRIL$_8$-TacF For | 5'- CAT ATA TCC AGA AGT ATC TTG CCC TGA GCT ATA TGG GTA TTC -3’ | 65.9 | This study |
| BRIL$_9$-TacF For | 5'- CAT ATA TCC AGA AGT ATC TTC TGA GCT ATA TGG GTA TTC - 3’ | 62.7 | This study |
| BRIL$_{10}$-TacF For | 5'- CAT ATA TCC AGA AGT ATC TTA GCT ATA TGG GTA TTC - 3’ | 59.7 | This study |
| BRIL-TacF Rev | 5'- AAG ATA CTT CTG GAT ATA TGC ATT GCG GGT-3’ | 60.6 | This study |
